# Ecological niche stability of *Biomphalaria* intermediate hosts for *Schistosoma mansoni* under extreme flooding and seasonal change

**DOI:** 10.1101/2025.01.07.631649

**Authors:** Melissa A. Iacovidou, Anatol M. Byaruhanga, Fred Besigye, Betty Nabatte, Narcis B. Kabatereine, Goylette F. Chami

## Abstract

Understanding spatiotemporal distributions and niches of vectors and intermediate hosts for ecologically dependent pathogens is crucial for identifying endemic areas, assessing habitat suitability for transmission, and targeting interventions for both the environment and humans. This study focuses on *Biomphalaria sudanica* and *B. stanleyi*, intermediate hosts of intestinal schistosomes, with over 700 million people at risk in endemic areas. We identified how extreme flooding and seasonal changes influence habitat preferences and species interactions across 674 sites in 52 villages in rural Uganda between 2022–2024. A comprehensive analysis of ground truth data was conducted, covering spatiotemporal information, site characteristics, physicochemical parameters, ecological factors, and human activities. Spatiotemporal models incorporating a new polygon-based method to account for space, bypassing limitations of administrative boundaries, with time as a fixed effect were developed to analyse snail abundance. *B. sudanica* was associated with marshy sites near lake shorelines and presence of hyacinths, while *B. stanleyi* was more likely found in deeper waters with *Vallisneria* plants. However, both species often cohabited at the same sites. The extent of habitat suitability for each species fluctuated seasonally and more starkly with extreme flooding resulting in switching of dominance between species. Our study shows that climatic variations may influence local changes in habitat suitability without necessitating an expansion of environmental areas. By providing a robust, generalisable spatiotemporal modelling pipeline, our study enables precise tracking of dynamic ecological niches in a changing climate that, if replicated in other areas, can be used to better target environmental and human interventions.

## Introduction

Schistosomiasis is a neglected tropical disease caused by *Schistosoma* trematode flukes [1] with over 250 million people infected globally, mostly in sub-Saharan Africa [2]. Infections can lead to lifelong morbidities, including irreversible fibrosis of the bladder, spleen, and liver depending on the schistosome species [1]. Transmission relies on freshwater snails acting as intermediate hosts. Yet, schistosomiasis control efforts primarily focus on mass drug administration (MDA) with implementations based on administrative units of endemic countries that neglect the complex spatiotemporal distributions of snails and their impact on infection prevalence. Recent World Health Organization guidelines strongly recommend environmental interventions in the form of water treatment and snail control [3]. Integrated approaches thus far have focused on precision mapping of snail habitats [4–6]. However, there remain outstanding questions about the ecological niches of different snail species, variability with climate change, and the best approaches for spatiotemporal modelling. Here we focus on a common genus of *Biomphalaria* in sub-Saharan Africa to assess habitat suitability in the context of seasonal change and extreme events such as flooding.

*Biomphalaria* snails act as intermediate hosts for *S. mansoni*, facilitating its asexual reproduction and the release of thousands of infectious cercariae from a single miracidium and single snail [1]. This amplification effect means that only a few hosts are required to infect entire communities, thereby complicating efforts to map and control infection risks, especially combined with the microgeographical focus of snail habitats that can vary on the basis of a few meters [7]. Snail distributions are made even more difficult to predict due to seasonal ecological variability such as rainfall, temperature, and changes in conditions specific to freshwater sites such as vegetation type and growth [8]. Climate change and extreme weather events can further destabilise these habitats, altering snail populations and shifting transmission dynamics [9–15]. However, it remains an open question as to how such destabilisation affects habitat suitability, including whether 1) ecological niches are simply expanding spatially with stable characteristics in a changing climate or 2) ecological niche concepts need to be revisited and there are alternatives to simple habitat expansion.

Several studies have concluded that different *Biomphalaria* species have preferred ecological niches, resulting in the dominance of different water sites by each species, although the exact characteristics measured and identified have been inconsistent. It has been found that certain species prefer habitats along the shoreline, such as *B. sudanica* and *B. pfeifferi*, whereas other species prefer deeper waters, such as *B. stanleyi* and *B. choanomphala* [8, 16]. Snail abundance has been linked with certain vegetation (e.g. *B. sudanica* with hyacinth plants and *B. stanleyi* with *Vallisneria* plants [8, 17, 18]) and physico-chemical water parameters, such as water pH and water temperature, though there is disagreement regarding positive or negative associations for the climate-sensitive water parameters [8, 16, 17, 19, 20]. For these proposed ecological niches, spatiotemporal models are lacking for studying the seasonal variations of different sites across years, especially for *B. stanleyi*, which has been demonstrated to be a highly competent intermediate host with the potential to spread beyond its geographically focal area of Lake Albert in Uganda [21]. Where *B. stanleyi* is present, it has been shown that severe outcomes of schistosomiasis, such as periportal fibrosis and portal hypertension, are more common than in areas where it is absent [22]. There is a need to model potentially more straight-forward effects, such as changes in water temperature and more complex understudied events like extreme flooding to understand the stability and relevance of ecological niches.

To investigate the relationships between environmental factors and snail populations, statistical analyses of malacological data often use methods such as simple tests for associations [8, 20, 23], Maximum Entropy (MaxEnt) models [12, 14, 24–26], and regression analyses (e.g. GLMMs) [4, 16, 17, 27, 28]. These approaches have advanced our understanding of ecological niches and snail distributions; however, spatial factors are often overlooked. In studies surveying water sites over multiple timepoints, it is frequently unclear how sites are matched across time. While smaller-scale studies with fewer sites over shorter periods can track individual sites more easily [16, 23, 25, 28], larger-scale, annual studies that are needed to understand climate change often face challenges in matching sites over time, especially during extreme weather events [14, 27]. Additionally, spatial data often are aggregated into larger units [4, 20, 27], leading to the loss of indispensable information about snail microhabitats and spatial patterns. In some cases, individual site points are treated as random effects [16, 17, 28], but this fine scale is not necessarily informative nor spatially relevant for identifying meaningful spatial variation in snail distributions. In the simplest case, models report residual spatial correlation without addressing it explicitly in the model setup [17]. New approaches are needed to account for space in models seeking to assess ecological niches or habitat suitability.

We propose a generalisable spatiotemporal modelling pipeline designed to analyse the presence and stability of distinct ecological niches for different snail species. We collected data from a total of 674 water sites from three districts in rural Uganda, across four time-points representing different weather conditions (flooded, dry, rainy, and dry following extended rains) from 2022–2024. Our aims were to determine whether *B. sudanica* and *B. stanleyi* have distinct ecological niches and to evaluate the stability of these niches across seasons and climatic change, particularly in the context of extreme flooding.

## Materials and methods

### Study context

The study was conducted within the prospective cohort SchistoTrack [29]. This community-based cohort was established in Buliisa and Pakwach in Western Uganda, and Mayuge in Eastern Uganda, starting with 38 villages in 2022 and expanding to 52 villages as of 2024. The three districts feature distinct ecologies as Buliisa is situated along Lake Albert, Pakwach along the outlet of the Nile River from Lake Albert, and Mayuge along Lake Victoria. The communities in these rural districts heavily rely on water sourced from these water bodies and thus come into frequent contact with them.

Malacology data is collected at least annually. Here we examined all timepoints from 2022 (study baseline) to 2024. There were four malacology surveys with three annual collections in Jan–Feb from 2022 to 2024 and an additional seasonal collection in Oct– Nov 2023. To avoid confusion between the annual and seasonal collections in 2023, we henceforth refer to them as ‘2023a’ and ‘2023b’, respectively. All timepoints exhibited distinct weather conditions and therefore represent four seasonal scenarios in our analysis: flooded season (2022), dry season (2023a), rainy season (2023b), and dry season following extended rains (2024). We refer to them as such throughout the analysis.

Surveys for data collection were created using Open Data Kit (ODK) Collect (versions 2022.4.2, 2023.2.4, and 2024.1) and data were entered in the ODK surveys on Android devices (software version Android 9 and 10). ODK Central (versions 2022.3.1, 2023.2.1, and 2024.2.1) was used for data management and quality control. Further data and statistical analyses were performed using R (version 4.2.1) [30].

### Water sites and covariates

Surveys were conducted by eight malacologists working in pairs, assisted by local auxiliary workers. Sites within the catchment of the study that were reported to be frequented by village inhabitants by the local assistants were identified. Information was collected for all water sites, irrespective of the presence of snails. The length of the sites was limited to 15 m to capture potential variations in snail abundance associated with microhabitats [7].

Thus, long beaches and shorelines were divided into multiple sites. Seasonal ponds, defined as water bodies that dry up in less than two months, were excluded from the surveys.

The information collected at each site included: GPS coordinates (taken while the surveyor was standing in the water); water depths (shallowest and deepest parts); human activities frequently performed there; number of boats present; site landform; water speed; occurrence of environmental modifications; ground substrate; types of vegetation, live-stock, and fish present; water turbidity, temperature, and pH; total dissolved solids; conductivity; and start time of snail sampling. These were chosen based on theoretical importance from the literature.

In addition, several photographs were taken of the water sites. During the collection period (2022–2024), the malacologists aimed to visit the same water sites based on the GPS coordinates and the previously taken photographs. Any additional water sites indicated by the local workers were visited as well. However, it proved challenging to match water sites over the years due to significant environmental changes and other obstacles (e.g. presence of wild animals). Therefore, we did not attempt to correspond the water sites over time.

A set of 44 covariates was assembled to be used in variable selection based on the data collected at the water sites. The covariates were split into five broader categories: spatiotemporal information, general site characteristics, physicochemical parameters, environmental and ecological attributes, and human activities. Detailed descriptions of covariates along with collection procedures can be found in S1 Text. The numerical variables were scaled to have a mean of zero and a standard deviation of one across the entire time frame using the scale function in R.

### Snail sampling and outcomes

Snail sampling was performed by the two malacologists for 30 minutes. The process involved 15 minutes of scooping followed by 15 minutes of hand-picking. Vegetation at the site was agitated and standard scoops were passed along it to collect snails. Forceps were also used to extract snails. Only snails with visible flesh in the shell were collected.

Snails were placed into labelled containers with vegetation and a damp cloth and transported to a temporary laboratory habitat within a few hours where species identification was completed the following morning. External morphological characteristics were assessed using a dissecting microscope (Olympus) along with the guide by the Danish Bilharziasis Laboratory [31]. Species of the genus *Biomphalaria* identified included *B. sudanica*, *B. stanleyi*, *B. pfeifferi*, and *B. choanomphala*. The total number of both alive and dead snails was recorded for each water site. Dead snails were confirmed by gently poking the body with forceps and observing the response. In our analysis, we considered the total number of collected snails, as dead snails died during transportation. Collected snails were sorted in containers by species and site in groups of five. The containers were placed under direct sunlight for a maximum of eight hours and checked every 30 minutes for cercariae shedding. Groups that were identified as shedding were split into individual snail containers and were further monitored to identify the shedding snails. The dissecting microscope was used to confirm whether the snails were shedding human cercariae. The number of shedding snails per site was recorded.

We focused on the two most common species in our study, *B. sudanica* and *B. stanleyi*, with their abundance (counts of snails at each site) as our primary outcomes. The purpose was to identify whether each species has a distinct ecological niche preference, i.e. whether it favours specific environmental and ecological conditions and has distinct determinants that support its presence and abundance in water habitats. This analysis extended to investigating species coexistence/cohabitation, where both species are present at a given water site. This was further extended to species dominance, defined as the species with the highest absolute count of snails within a site, further examining whether dominance was stable or switched over seasons. The other two species commonly found in Uganda, *B. pfeifferi* and *B. choanomphala*, were not modelled due to their low numbers. We believe *B. pfeifferi* was present due to extreme flooding, which displaced them from their usual seasonal pond habitats. Furthermore, shedding snails were also only discussed descriptively due to their low numbers, and because the focus of this paper is on ecological niches related to presence and abundance.

### Spatial analysis

To understand the extent to which the snail counts of each of the two species at a given water site were similar or dissimilar to the counts in nearby locations we calculated spatial autocorrelation for each outcome. Using the spdep package [32], we first identified neighbouring observations by applying the dnearneigh function, setting a distance threshold equal to the maximum distance between any two sites within the same district. This ensured that sites from different districts were not considered as neighbours. We then computed Moran’s *I* [33], which quantifies spatial autocorrelation, using the moran.test function.

In the event of spatial autocorrelation being present, we constructed spatial units in the form of polygons that included the water sites, which were static to compare across timepoints. Despite the water sites being assigned to a village, they were not necessarily geographically located in that village. To accurately reflect this, we considered the households involved in the study and created a tessellation of polygons for each district separately based on the location of the households. Two approaches were explored for generating these polygons: (a) household clustering to capture spatial groupings independent of village boundaries (cluster-based polygons), and (b) household grouping based on self-reported village affiliations (village polygons). The cluster-based polygons were chosen for subsequent statistical modelling as they better aligned with the spatial focus of the study. Details of the alternative village-based approach are provided in S2 Text.

Constructing polygons based on household clustering was purely spatial and did not take into consideration the villages as defined by local residents or by the national government. To do this, we first applied a clustering algorithm to identify household clusters. Hierarchical clustering was chosen as it does not require any predefined number of clusters and does not allow for any points (households) to be treated as noise [34, 35], unlike density-based clustering [36]. A distance matrix between all the households in the district was calculated using the ‘Vincenty’ (ellipsoid) method [37]. To determine the optimal number of clusters (*k*), the gap statistic was considered as a goodness of clustering measure [38]. The gap statistic compares the within-cluster variation for different values of *k* in the actual data to that of a null reference distribution, which assumes no cluster structure. The optimal number of clusters is the *k* that maximises the gap statistic, indicating that the clustering structure in the data is significantly better than what would be expected by random chance. Using the factoextra package [39], we computed the gap statistic (with 500 simulations) and applied a hierarchical clustering algorithm with both the complete and average linkage methods [40–42], which ultimately gave us the same clusters. The polygons were constructed using the household locations; Voronoi polygons were generated for each household [43], and subsequently, these polygons were merged according to the cluster they belonged in. We identified which polygon each water site lay within for each timepoint and created the polygons variable.

For the construction of the tessellations we used the following packages: deldir [44], sp [45, 46], sf [47], and maptools [48].

### Statistical modelling

We fitted generalised linear mixed models (GLMMs) [49] with a zero-inflated negative binomial (ZINB) distribution to the snail counts for *B. sudanica* and *B. stanleyi* using the glmmTMB package in R [50]. A log link function was used for the count component to ensure no negative values are predicted and a logit link function for the zero-inflation component to ensure the probability of an excess zero is between zero and one.

While the distribution of counts is often modelled with a negative binomial distribution, we accounted for excess zeros by incorporating zero inflation, which considers structural zeros in addition to the zeros arising from the count process [51]. For instance, zeros in our snail counts may arise from the true absence of snails due to unsuitable habitats or other environmental conditions (structural/true zeros), measurement error such as the inability to collect snails on a given day (sampling/false zeros), or sampling variability, e.g. snails could have been present but perhaps due to microenvironmental changes on the day of sampling they are not (random/true zeros). Odds ratios (ORs) from the zero-inflation component of these models represent the odds of an observation being a structural zero. Thus, an OR *>* 1 indicates higher odds of snail absence, while an OR *<* 1 indicates lower odds. This contrasts with traditional logistic regression, where ORs reflect the odds of presence.

We constructed a minimally adjusted model including the timepoint and district covariates as fixed effects and the polygons covariate as a random effect (in both the conditional and zero-inflation parts of the GLMM). We then performed variable selection on the remaining 41 variables (described in S1 Text) as outlined below.

### Variable selection

We performed variable selection separately for each species to determine the remaining fixed effects. First, we selected variables for the zero-inflation part of the model by adding them one at a time and performing a likelihood ratio test (LRT) to assess their significance [52]. We considered variables with a *p*-value *<* 0.05 to be significant. We updated the zero-inflation part of the minimally adjusted model to include the chosen variables. Next, we included one variable at a time in the conditional part of the updated model and performed LRTs to identify significant variables. This process resulted in two GLMMs, one for each species. *B. stanleyi* snails are not found in Lake Victoria, therefore the district of Mayuge was excluded from the analysis for this species. Variance inflation factors (VIFs) were used to investigate multicollinearity in the fully adjusted models [53] using the multicollinearity function from the performance package [54]. We removed variables (that were not part of the minimally adjusted model) with VIF *>* 5 one by one until no multicollinearity was detected, with a maximum of three variables ultimately being removed. To enable direct interpretation of the model coefficients, we calculated odds ratios (ORs) for the zero-inflation part of the model and rate ratios (RRs) for the conditional part by exponentiating the coefficient estimates.

### Model validation

The DHARMa package was used to perform goodness-of-fit tests on simulated residuals [55]. We measured spatial autocorrelation in the residuals of the fully adjusted models using Moran’s *I*, as before, to determine whether any unaccounted-for autocorrelation remained. Additionally, we computed the corrected Akaike information criterion (AICc) for the minimally and fully adjusted models to estimate the relative quality of each.

We applied multiple cross-validation methods to evaluate the predictive capacity of the models and answer key questions about generalisability in different spatial and temporal scenarios, particularly in Uganda. We conducted stratified 5-fold cross-validation, where we randomly sampled from each timepoint, district, and polygon. Sampling from each timepoint ensured that the temporal structure was preserved in each fold, accounting for the distinct climate profiles associated with each timepoint, which could otherwise introduce bias. Similarly, spatial sampling within each district ensured that the spatial structure at the district level was maintained, reflecting the ecological differences influencing snail habitats and biodiversity, shaped by the different water bodies each district lies along. Furthermore, sampling within each polygon enabled local variations, that may not necessarily be district-wide, to be taken into account. These stratified 5-fold approaches allowed us to answer how well the models generalise and perform across space and time. To assess sensitivity and robustness, we also performed leave-one-out cross-validation (LOOCV), where we trained on all polygons except one, which was used as the test dataset. This helped assess whether the models were sensitive to specific polygons and understand how the models perform at the polygon level. We calculated the receiver operating characteristic (ROC) for the probability of a site having zero snails and computed the area under the curve (AUC) value for each method. The caret package was used for model training and cross-validation [56], and the pROC package to compute the ROC and AUC values [57].

### Modelling pipeline

A summary of the methods and generalisable spatiotemporal modelling pipeline—with a focus on key decision points—is presented in Figure 1.

**Figure 1:**
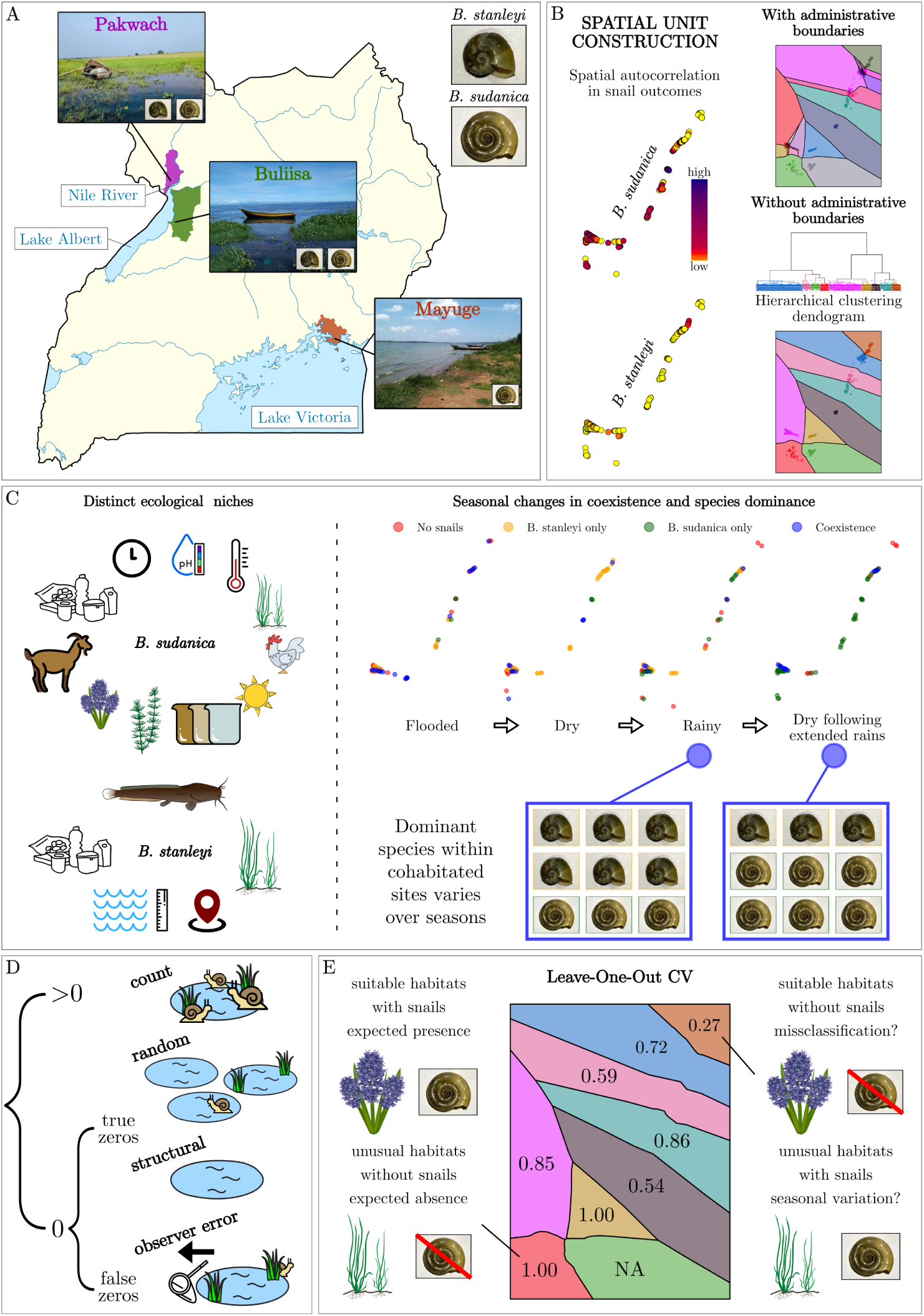
Methodological pipeline. (**A**) Study context showing examples of sites at each district and the main *Biomphalaria* snails studied. (**B**) Spatial unit construction and spatial autocorrelation in outcomes. (**C**) Ecological niches (left) and seasonal variations in coexistence and dominance (right) with examples of dominance in cohabitated sites. (**D**) Reasons and differences between true and false zeros. (**E**) Model validation with example for high-and low-performing polygons using LOOCV.

## Results

### Summary of water sites and snails

We observed a total of 674 water sites across four timepoints from 2022 to 2024. During the initial flooded season (2022), we collected data from 143 sites across 38 villages, with two villages reporting no water sites. In the subsequent timepoints, the number of study villages increased to 52, resulting in a corresponding increase in the number of sites. Specifically, we surveyed 177 sites during the dry season (2023a), with seven villages reporting no water sites; 175 sites during the rainy season (2023b), with five villages reporting no sites; and 179 sites during the dry season following extended rains (2024), with five villages again reporting no sites. Four of the newer villages were consistently site-free. When focusing only on the initial 38 villages, the number of observed sites was highest during the flooded season (2022) with 143 sites and decreased across the timepoints, with 138, 134, and 141 sites recorded in 2023a, 2023b, and 2024, respectively. Table 1 shows a breakdown of sites by timepoint and district.

**Table 1:** Breakdown of number of sites per district and timepoint.

|  | **2022**<br>(Flooded) | **2023a**<br>(Dry) | **2023b**<br>(Rainy) | **2024**<br>(Dry following<br>extended rains) | **Total** |
| --- | --- | --- | --- | --- | --- |
| **Buliisa** | 59<br>(41.3%) | 61<br>(34.5%) | 64<br>(36.6%) | 68<br>(38.0%) | 252<br>(37.4%) |
| **Pakwach** | 40<br>(28.0%) | 66<br>(35.0%) | 62<br>(35.4%) | 62<br>(34.6%) | 230<br>(34.1%) |
| **Mayuge** | 44<br>(30.8%) | 50<br>(28.2%) | 49<br>(28.0%) | 49<br>(27.4%) | 192<br>(28.5%) |
| **Total** | 143 | 177 | 175 | 179 | **674** |

Over all timepoints and sites, a total of 61,459 snails were collected, with 62.7% (38505/61459) being *B. sudanica*, 31.8% (19516/61459) *B. stanleyi*, 3.7% (2253/61459) *B. pfeifferi*, and 1.9% (1185/61459) *B. choanomphala*. Furthermore, a total of 618 snails were observed to be shedding human cercariae, out of which 20.7% (128/618) were *B. sudanica*, 75.1% (464/618) *B. stanleyi*, 2.9% (18/618) *B. pfeifferi*, and 1.3% (8/618) *B. choanomphala*. Breakdowns by district and timepoints for the number of collected snails and shedding snails are presented in S3 Tables and S4 Figures.

### Spatial and seasonal variation in covariates

We present descriptive statistics of covariates exhibiting significant differences over districts and timepoints in S5 Tables. Numerous covariates had significant differences over districts or timepoints or both, highlighting the spatiotemporal variability present in our data. Plots of distributions of key covariates including altitude, water depth, water pH, water temperature, conductivity, and landform are shown in S6 Figures.

### Spatial autocorrelation and spatial unit

Calculating Moran’s *I* for the two snail outcomes, we observed the presence of spatial autocorrelation prior to model building. Specifically, *I* = 0.131 (*p*-value *<* 0.001) for *B. sudanica* and *I* = 0.010 (*p*-value *<* 0.001) for *B. stanleyi*. Cluster and village-polygons were compared. For the polygons reliant on household clustering, the optimal number of clusters, *k*, for each district was computed as 9, 15, and 12 for Buliisa, Pakwach, and Mayuge, respectively (S7 Figure). The number of spatial units needed was reduced by 30.8% when moving from self-reported administrative villages (52) to cluster-based polygons (36). The resulting district tessellations including the water sites are visualised in Figure 2. Polygons based on administrative village units provided less reliable structures requiring manual, subjective reassignment of households to village polygons (S8 Figures). Thus, cluster-based polygons were used in all models as a random effect to account for spatial autocorrelation.

**Figure 2:**
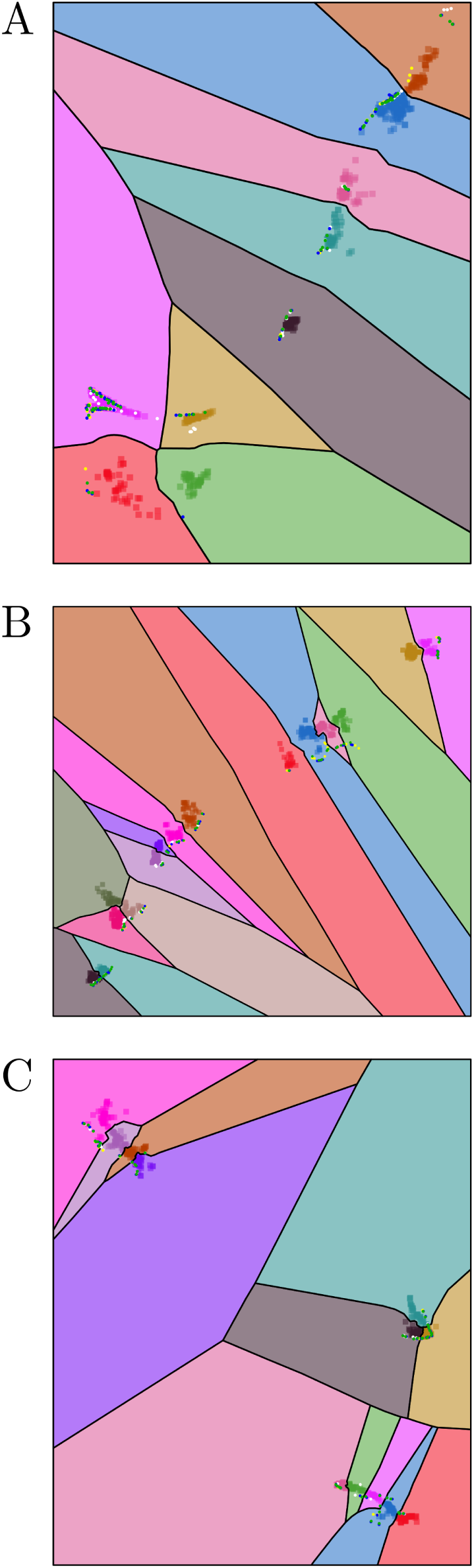
District tessellations including water sites. A. Buliisa, B. Pakwach, C. Mayuge. Squares represent individual households and dots represent water sites. The site colours correspond to the four timepoints: white – 2022; yellow – 2023a; blue – 2023b; green – 2024.

### Ecological niche of *B. sudanica*

We observed the presence of *B. sudanica* in 63.8% (430/674) of all water sites over time (S9 Figure). Figure 3 presents the OR and RR plots of the fully adjusted model for *B. sudanica*, and the model without Mayuge is shown in S10 Figure.

**Figure 3:**
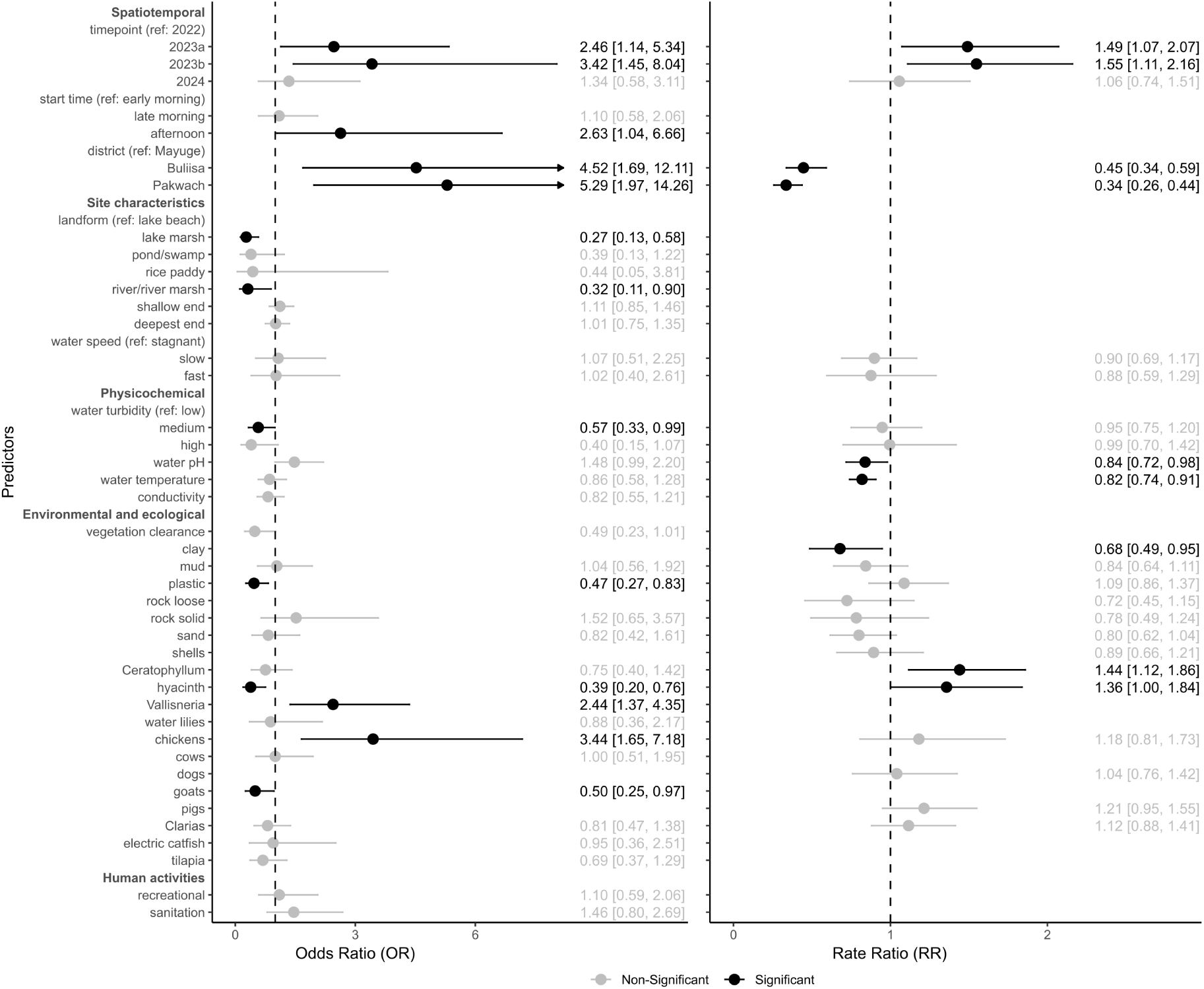
Fully adjusted *B. sudanica* model. Odds Ratios (left) and Rate Ratios (right) for the zero-inflation and conditional components of the model, respectively. Dots correspond to the exponentiated coefficient estimates and lines correspond to the 95% confidence intervals. Arrows indicate that the confidence interval continues off the plot.

Water sites in Buliisa and Pakwach had 4.63 (95% CI: 1.72 to 12.47) and 5.27 (95% CI: 1.96 to 14.11) times higher odds of having excess zeros compared to sites in Mayuge. Excess zeros refer to sites with no observed snails, which may result from unfavourable ecological conditions or sampling challenges. Seasonal variation in excess zeros was observed, as sites in the 2023a (dry) and 2023b (rainy) timepoints were more likely to have excess zeros compared to 2022 (flooded) (OR = 2.23, 95% CI: 1.02 to 4.81 and OR = 2.98, 95% CI: 1.26 to 6.95, respectively). Collecting snails in the afternoon had 2.67 (95% CI: 1.06 to 6.60) times higher odds of leading to excess zeros compared to early morning collection. The most common site landform was lake beach and was therefore used as the reference category. Lake sites with a marshy landform had a 72% lower chance of having excess zeros compared to a beachy landform (OR = 0.28, 95% CI: 0.13 to 0.59), while no significant differences in excess zeros were found for other landforms. Several ecological factors also influenced the presence or absence of snails. The presence of hyacinth was associated with a reduced likelihood (OR = 0.40, 95% CI: 0.21 to 0.79) of sites having excess zeros, while the presence of Vallisneria plants led to 2.60 (95% CI: 1.45 to 4.65) times higher odds of observing excess zeros compared to when no Vallisneria plants were present. Additionally, the odds of observing excess zeros were 3.47 (95% CI: 1.64 to 7.26) times higher when chickens were present compared to when they were not. Conversely, the presence of goats reduced the likelihood of sites having excess zeros (OR = 0.48, 95% CI: 0.25 to 0.93). Similarly, sites with plastic and other waste had a reduced likelihood of excess zeros (OR = 0.46, 95% CI: 0.26 to 0.81) as compared to the absence of these ground substrates. Medium turbidity was associated with a reduced likelihood of excess zeros (OR = 0.55, 94% CI: 0.31 to 0.96) compared to low turbidity.

The expected snail abundance was lower in the Western Districts of Buliisa (RR = 0.45, 95% CI: 0.34 to 0.59) and Pakwach (RR = 0.34, 95% CI: 0.26 to 0.44) when compared to the Eastern District of Mayuge (right section of Figure 4). Concerning the changes over time, the expected count was lower during the extreme flooding in 2022, with counts in 2023a (dry) and 2023b (rainy) being 1.48 (95% CI: 1.07 to 2.06) and 1.55 (95% CI: 1.11 to 2.16) times higher, respectively, while counts in 2024 (dry following extended rains) showed no significant difference. The presence of *Ceratophyllum* and hyacinth plants led to a higher number of snails on average than those without these plants (RR = 1.44, 95% CI: 1.11 to 1.85 and RR = 1.36, 95% CI: 1.00 to 1.84, respectively), though hyacinth plants were borderline significant (*p*-value = 0.048). The presence of any clay as ground substrate led to a lower snail abundance (RR = 0.67, 95% CI: 0.48 to 0.94). Physicochemical water parameters associated with snail abundance included water pH where for each one standard deviation (*≈* 0.8 pH units) increase the expected number of snails declined by 16% (RR = 0.84, 95% CI: 0.72 to 0.98). Similarly, a one standard deviation increase in water temperature (*≈* 1.4 ^◦^C) was associated with a decrease in average abundance by 18% (RR = 0.82, 95% CI: 0.74 to 0.91).

**Figure 4:**
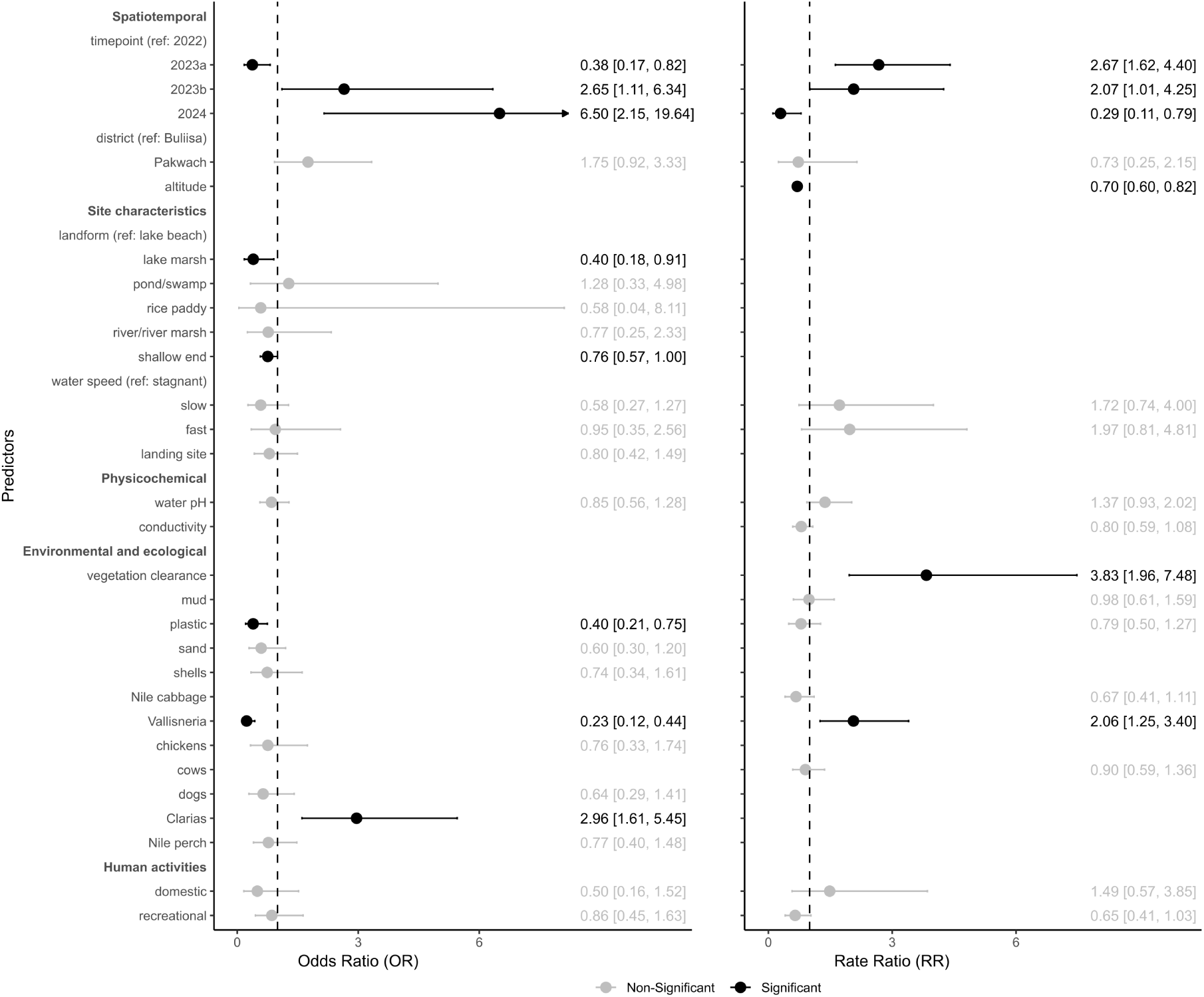
Fully adjusted *B. stanleyi* model. Odds Ratios (left) and Rate Ratios (right) for the zero-inflation and conditional components of the model, respectively. Dots correspond to the exponentiated coefficient estimates and lines correspond to the 95% confidence intervals. Arrows indicate that the confidence interval continues off the plot.

### Ecological niche of *B. stanleyi*

*B. stanleyi* snails were present in 40.7% (196/482) of the observations, excluding water sites in Mayuge (S11 Figure). Figure 4 presents the OR and RR plots of the fully adjusted model for *B. stanleyi*.

Comparing the likelihood of sites in Buliisa and Pakwach exhibiting excess zeros, indicating the absence of snails, there was no significant difference between the districts (*p*-value = 0.085). Seasonal variation in excess zeros was noticed; compared to the flooded (2022) timepoint, the odds of observing excess zeros were 2.65 (95% CI: 1.11 to 6.34) times higher in the rainy season (2023b) and 6.50 (95% CI: 2.15 to 19.64) times higher in the dry season following extended rains (2024), while the odds of a site having excess zeros decreased by 62% (OR = 0.38, 95% CI: 0.17 to 0.82) in the dry season (2023a). Lake sites with a marshy landform had 60% lower chance of having excess zeros present compared to lake sites with a beachy landform (OR = 0.40, 95% CI: 0.18 to 0.91). The presence of any *Clarias* fish was associated with a higher likelihood of excess zeros (OR = 2.96, 95% CI: 1.61 to 5.45). Vegetation type and presence of plastic or other waste at sites were correlated with the decreased likelihood of finding excess zeros. The odds of observing excess zeros were approximately 77% lower when *Vallisneria* plants were present compared to their absence (OR = 0.23, 95% CI: 0.12 to 0.44) and 60% lower when plastic or other waste was present (OR = 0.40, 95% CI: 0.21 to 0.75, respectively). One standard deviation (*≈* 12.1 cm) increase in water depth for the shallow end of the site led to a 25% decrease in the odds of excess zeros (OR = 0.76, 95% CI: 0.57 to 1.00).

There was no significant difference between the expected number of snails between the two districts (*p*-value = 0.562). However, differences were observed between seasons, where the expected number of snails was 2.67 (95% CI: 1.62 to 4.40) times higher in the dry season (2023a) and 2.07 (95% CI: 1.01 to 4.25) times higher in the rainy season (2023b), compared to the flooded season (2022). On the other hand, the dry season following extended rains (2024) exhibited a 71% (RR = 0.29, 95% CI: 0.11 to 0.79) decrease in the expected number of snails compared to the flooded season (2022). The presence of *Vallisneria* plants was associated with an expected snail abundance that was 2.06 (95% CI: 1.25 to 3.40) times higher compared to sites without *Vallisneria* plants. Furthermore, clearance of vegetation also led to the expected number of snails being 3.83 (95% CI: 1.96 to 7.48) times higher than sites where this had not occurred. One standard deviation (*≈* 41 m) increase in altitude was associated with a 30% lower snail abundance (RR 0.70, 95% CI: 0.60 to 0.82).

### Species cohabitation and dominance switching

The two species had many overlapping ecological dimensions, yet the ecological niches were distinct. For the zero-inflated component, there was overlap in selected variables including landform, shallow end depth, water speed, water pH, plastic and sand ground substrates, *Vallisneria* plants, chickens, *Clarias* fish, and recreational activities, whereas only water speed, water pH, mud and plastic substrates overlapped for the conditional components. From these variables, only the coefficients of plastic and *Vallisneria* were significant for both models (in the zero-inflation part) with the effect of *Vallisneria* in opposite directions for each species.

Despite distinct ecological niches, we detected the coexistence of the two species in 17.2% (83/482) of all sites across time (excluding Mayuge) with the coexistence peaking in the flooded season at 30.3% (30/99) in 2022 then varying seasonally from 17.3% (22/127) in 2023a, 10.3% (13/126) in 2023b, and 13.8% (18/130) in 2024. Sites with coexistence were largely clustered together within a few polygons. The spatiotemporal variations of the sites with no snails, *B. stanleyi* only, *B. sudanica* only, and coexistence of both species are shown in panels A and B in Figure 5.

**Figure 5:**
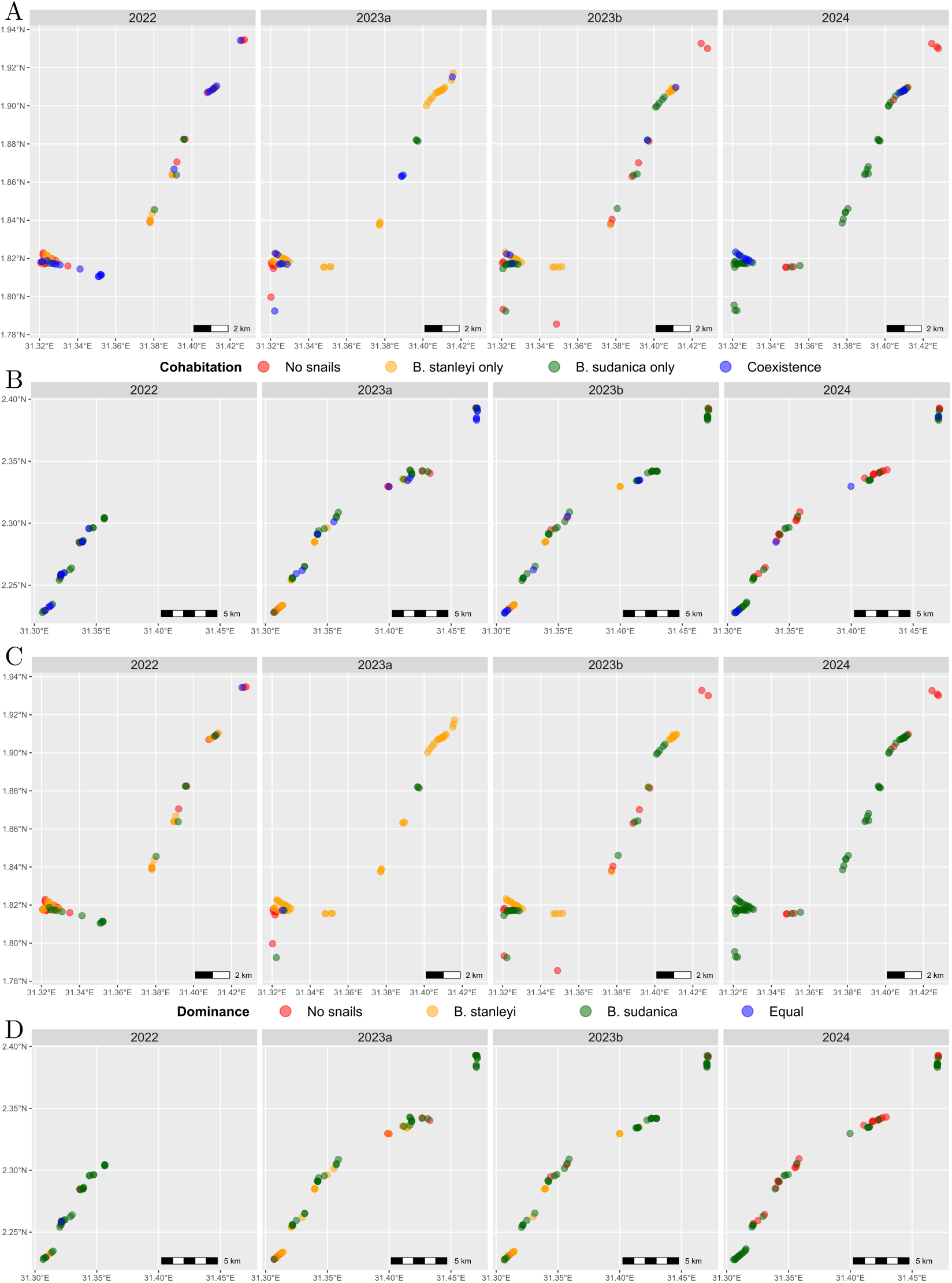
Cohabitation and within-site dominance of snail species. Cohabitation: A. Buliisa, B. Pakwach. Dominance: A. Buliisa, B. Pakwach.

As an alternative to the strict measure of absence of one species to indicate the lack of coexistence, we also investigated dominance (differences in absolute abundance), to understand potential dominance switching depending on the season. Over 70% of the cohabitated sites (59/83) had an absolute difference between species of more than 20 snails (median 51, IQR 131). Overall, 51.2% (247/482) of sites were dominated by *B. sudanica* and 29.5% (142/482) by *B. stanleyi* with the remaining sites having no snails or the same number of snails from both species (panels C and D Figure 5). The dominant species in most sites between the two districts was not the same across seasons. In Buliisa the switches between dominance were more evident, with somewhat balanced dominance (in terms of how many sites each species is dominant at) in the flooded and rainy seasons where, in contrast, there was a clear single species dominance in the two drier timepoints.

### Model validation

Results from goodness-of-fit tests on simulated residuals are shown in S10 Figure. There was no remaining spatial autocorrelation in either model as calculated by Moran’s *I* which was close to zero for both (*−*0.004, *p*-value = 0.8 and *−*0.003, *p*-value = 0.6 for the residuals of the *B. sudanica* and *B. stanleyi* models, respectively). The AICc was calculated for the minimally and fully adjusted models; for *B. sudanica* the AICcs were 5426 and 5295, and for *B. stanleyi* 2602 and 2489. The selected predictors improved the fit of the minimally adjusted models, which already included the key spatiotemporal features of timepoint, district, and polygons, for both species.

The AUC values computed for the cross-validation methods were based on the probability of structural zeros (absence of snails) predicted by the models. The average AUC values for the stratified 5-fold cross-validation, when we sampled from within each polygon, district, and timepoint, were 0.80, 0.81, and 0.81 respectively for *B. sudanica* and 0.79, 0.81, and 0.83 for *B. stanleyi*. For the leave-one-out cross-validation (LOOCV), AUC values ranged from 0.125 to 1, with a median of 0.67 for *B. sudanica* and from 0.20 to 0.96, with a median of 0.75 for *B. stanleyi*. An observed pattern for the polygons with low predictive capacity for *B. sudanica* was the absence of snails when hyacinth, goats, and/or plastic (waste) were present. These variables were significant in the zero-inflation part of the model and thus the predictive capacity of the model was low in polygons with sites having these characteristics. Similarly, for *B. stanleyi*, absence of snails when *Vallisneria* plants and/or plastic were present as well as presence of snails when *Clarias* fish were present led to a lower predictive performance.

## Discussion

Extreme events or seasonal trends drive schistosome transmission dynamics and influence habitat suitability. Understanding these spatiotemporal influences is essential for identifying areas of ongoing transmission and adapting interventions to climate change. Here we focused on the spatiotemporal distributions and ecological niches of *B. sudanica* and *B. stanleyi* snails, which are intermediate hosts of *S. mansoni*, in schistosome endemic areas of rural Uganda. We surveyed 674 water sites from the years 2022–2024 across four ecologically distinct sampling periods. Our study revealed that the distributions of *B. sudanica* and *B. stanleyi* were strongly influenced by extreme flooding and seasonal variations, leading to species cohabitation, dominance switching, and instability in the distinction of their ecological niches.

The ecological niche of *B. sudanica* found here was influenced by site characteristics, ecological factors, and physicochemical water parameters. These snails were found in sites with medium or high water turbidity, as opposed to low turbidity. *B. sudanica* snails were more likely in the presence of plastic and other waste, as well as goats, whereas the presence of chickens and *Vallisneria* plants created unfavourable conditions. Perhaps goats enhanced the habitat by providing food through their excreta whereas chickens could have created a predatory environment. Snails were less likely to thrive in sites with clay substrates and were found more often in lower water temperatures. In this case, our findings might suggest that increasing water temperature due to climate change, though suggested to widely support schistosome transmission elsewhere [10], can be detrimental to certain snail species, such as *B. sudanica*. By accounting for a wide range of variables, our analysis also helps address the inconsistent relationships of water pH found in the literature, suggesting both a positive and negative association for *B. sudanica* [8, 16, 19, 58]. Importantly, during collections conducted in the afternoon compared to early morning, we found that snail absence was increased, supporting that afternoons may be a less risky time for human transmission due to lower cercarial shedding by snails [59]. A positive association with *Ceratophyllum* and hyacinth plants, as well as preference for marshy (over beachy) sites confirms what has been seen in malacological studies [8, 17, 20, 60–62].

The ecological niche of *B. stanleyi* was influenced by site characteristics and ecological factors. The presence of *Clarias* fish was unfavourable to *B. stanleyi* snails, suggesting that they might prey on them. Presence of plastic and other waste seemed to be favourable, possibly suggesting that waste materials provide microhabitats or resources that snails take advantage of, perhaps due to the potential implications of waste on water quality, although the specific mechanisms remain unclear. Water depth also influenced snail distribution, with deeper shallow ends proving preferential. A significant relationship with altitude was observed, with higher elevations being unfavourable for *B. stanleyi* —perhaps an indicator for air temperature as lower altitudes are associated with higher temperatures. This may explain its absence in Mayuge, where the average altitude (1148 m) is significantly higher than in Buliisa (621 m) and Pakwach (625 m). Differences within districts should be taken with caution as vertical measurements from GPS devices are prone to error. The positive association of *B. stanleyi* with the presence of *Vallisneria* plants was reconfirmed, but we found *B. stanleyi* was more common in marshy landforms than beaches, challenging the sandy habitat preference observed elsewhere [8].

The two species were observed to coexist across seasons, with their seemingly distinct ecological niches reflecting flexible habitat preferences rather than fixed boundaries. Studies often portray habitat differentiation, rarely exploring the conditions favourable across species (such as plastic or other waste and vegetation clearance) or even the possibility of cohabitation [8, 20]. However, our results not only show that the species do coexist, they also indicate that the species dominance at a cohabitated site can shift over time, underscoring the dynamic role of seasonal variations in shaping their ecological interactions. Clear patterns of seasonal dominance switching were particularly evident in the district of Buliisa, where *B. stanleyi* dominated during the dry season and *B. sudanica* became dominant during the dry following extended rains season. In the flooded and rainy seasons, we saw a balance between species, blurring habitat distinctions, with highly clustered cohabitated sites. During the dry and rainy seasons, we found that sites were more likely to have no *B. sudanica* present, however, when present at sites, the abundance was higher compared to the other seasons, suggesting that while many sites became unsuitable, those that remained suitable supported larger populations. This pattern implies that conditions during the dry and rainy seasons may reduce overall habitat availability but create temporary niches for *B. sudanica* in the remaining habitats. This supports previous studies where normal water levels were more favourable than flooded conditions [8, 16, 19]. In contrast, *B. stanleyi* habitat suitability increased (as per higher abundance) during the dry season, compared to the flooded season, consistent with the literature [8]. During the rainy season, habitat suitability declined, but the remaining suitable sites supported larger snail populations. Extreme flooding, characterised by high water levels, may reduce light penetration to submerged vegetation, decreasing oxygen concentrations and influencing snail dynamics. These conditions can be exacerbated by decomposing plants such as *Vallisneria* and hyacinths, which are otherwise beneficial to *B. stanleyi* and *B. sudanica*, respectively. However, under flooded conditions they might create harmful environments for snails over time, highlighting the complex interaction between vegetation presence, water quality, and seasonal effects shaping habitat suitability and snail abundance over time. Even established knowledge of species’ climatic preferences, once compared across species provides evidence of otherwise overlooked shared snail habitats.

Environmental modification, such as vegetation clearance, has been proposed as a sustainable intervention to control snails and reduce human infection risk. However, our findings indicate that vegetation clearance had a borderline significant effect on the presence of *B. sudanica* and increased the abundance of *B. stanleyi*. These results suggest that disturbances to vegetation structure or water conditions can sometimes create favourable environments for snails, contradicting previous studies that linked submerged vegetation removal to reduced *Biomphalaria* abundance [63, 64]. This study, conducted in West Africa along the Senegal River, focused on *B. pfeifferi* and *Bulinus* snails, as well as *Ceratophyllum* plants, which suggests that the findings may not be generalisable to other species, regions, or types of vegetation. Therefore, vegetation removal could yield mixed results, depending on the species and ecological context. This raises the question: Can snail control interventions designed for one species be effective across multiple species, or would it require a detailed, species-specific strategy? The complexity increases when aiming for a multi-species approach, as different species would have different ecological niches that need targeting, leading to infeasible or ineffective strategies. Additionally, interventions like environmental modifications may have unpredictable or mixed effects, as the relationship between vegetation structure, water quality, and snail populations is often complex and context dependent. As such, a more targeted and integrated approach, possibly incorporating multiple intervention strategies, may be necessary to address the challenges of snail control, and longer-term efforts to offset climate change cannot be ignored. Future research at a large scale is still needed to investigate the generalisability of environmental modifications across different species, regions, and types of vegetation.

*B. stanleyi* snails have been understudied for not only their interactions with co-endemic snail species, but importantly also human transmission. *B. stanleyi* snails are native to Lake Albert and closely related to *B. pfeifferi* [65]. While they have been previously reported in Lake Chad and Lake Cohoha [66], their current presence in these regions remains unclear. Previous studies have explored aspects of their phylogeny [67], morphology [68], ecology [8], and population dynamics [69], yet *B. stanleyi* is largely overlooked compared to other *S. mansoni* intermediate hosts such as *B. sudanica*, *B. pfeifferi*, and *B. choanomphala*, which are more widespread. This underrepresentation is particularly concerning given that areas around Lake Albert are characterised by a high burden of life-threatening morbidity related to *S. mansoni*, such as portal hypertension and high infection prevalence [22, 70], raising questions about the role of *B. stanleyi* in parasite virulence and transmission. Experimental investigations have demonstrated that *B. stanleyi* can act as an efficient host for the *S. mansoni* isolates from Lake Victoria [21], performing even better than native Lake Victoria snails. If this species were to migrate to other regions, such as Lake Victoria or even worse cross borders to other countries, it could impact transmission dynamics and infection rates, potentially undermining ongoing control efforts, such as MDA. Further research is necessary to understand the potential migration of this species into other *S. mansoni*-endemic regions and its implications for schistosomiasis control.

The spatial unit used for environmentally mediated pathogens, such as schistosomes, affects how we understand transmission and the effectiveness of interventions. We developed new methods to account for spatial correlation in environmentally dependent models of infectious diseases with intermediate hosts or vectors that have limited mobility ranges. Especially for schistosomiasis, the need for a more relevant spatial unit has been evident and overlooked, with districts still acting as the implementation unit of interventions in many countries [71, 72]. Flooding or dryness causes shorelines to move, leading to uncertainty as to which water sites are the same over time, especially in areas with extreme weather variations. By constructing spatial units based on households rather than water sites, we created a more stable and robust framework for tracking water site locations over time. This approach minimises the risk of changes between surveys, making it a reliable tool for future work in identifying focal points of transmission. Additionally, it offers flexibility in linking with administrative boundaries if needed, as well as improving the relevance of geographical clustering for interventions. The effectiveness of our method for dealing with spatial autocorrelation was evident when initial autocorrelation present in the outcomes was no longer present in the residuals of our models that incorporated the polygons as a random effect. The polygons can also be used in future work investigating human infection and exposure and infection models, especially meta-population and individual-based models.

We validated our models to assess the generalisability for spatial and seasonal scenarios to inform potential use cases. The consistent results obtained from the stratified 5-fold cross-validation for both species sampling from each district, polygon, and timepoint (a conventional internal validation method [73, 74]) show the models were robust across the spatial and temporal groups tested with high predictive capacity (*>* 0.80 AUC). However, looking at the LOOCV results, we noticed that there were some failure cases (with low AUC values), suggesting that the models were more locally relevant for a subset of geographical areas. We found that the variables with significant coefficients in the zero-inflation component of the models played a role in the failure cases. For example, strong relationships were seen in *B. sudanica* presence and hyacinth plants and *B. sudanica* absence and *Vallisneria* plants, leading to low-performing polygons when sites have opposite conditions. This also could be the case of misclassification of zeros, for example with false zeros being considered as structural zeros. Such failure is to be expected in select polygons and should be more widely reported in future studies applying statistical models. These cases underscore the challenges of predicting species presence and abundance in highly variable ecological landscapes, even within the same districts.

In an ideal scenario, the identification of false zeros would be straightforward, allowing for their removal or direct modelling from the data. A limitation in our spatial approach is that it requires a spatial boundary to be defined, which in this case was the study catchment area to enable ground truth data collection. As a result, water sites could potentially be missing within the catchment of our tessellations. Another limitation arises from the treatment of districts and timepoints as fixed effects, which prevents the model from predicting unseen districts or timepoints as a whole, but can be overcome in future studies applying our pipeline if there are a large number of districts or timepoints and each are incorporated as random effects.

## Conclusion

In this paper, we challenge the hypothesis that *B. sudanica* and *B. stanleyi* have distinct, stable ecological niches. We provide evidence that cohabitation is not uncommon and driven by environmental and seasonal disturbances. Shifts in species dominance across timepoints highlight the importance of considering extreme weather events and their impacts on schistosome transmission, particularly in a time where climate change is impacting infectious disease dynamics worldwide. The variability in snail presence and abundance across different timepoints and districts, influenced by both natural and anthropogenic factors, suggests that targeted control measures should be adapted based on seasonal and ecological conditions. Monitoring aquatic vegetation and physicochemical water parameters, consistently linked to abundance, should be prioritised, along with surveying during different weather conditions to identify nuanced microhabitats that might be created. The polygon-based spatial approach used in this study proved effective in capturing variability across flexible, ecologically relevant units. Our spatial approach may enable future studies to integrate snail distribution with human mobility and infection data to refine force-of-infection models and optimise control strategies. If extended to other contexts and species, our findings and methods can inform locally relevant models estimating environmental suitability of schistosomes in a changing climate.

## Ethics approval

Data collection and use were reviewed and approved by Oxford Tropical Research Ethics Committee (OxTREC 509-21), Vector Control Division Research Ethics Committee of the Uganda Ministry of Health (VCDREC146), and Uganda National Council of Science and Technology (UNCST HS 1664ES).

## Data and code availability

All relevant metadata are provided within the manuscript and supplementary material and code to reproduce the pipeline is available in a GitHub repository.

## Acknowledgements

We thank the SchistoTrack group for valuable feedback and insights during group meetings and conversations. We thank all field teams, especially the malacologists, the local auxiliary workers, and village health team members.

## Author contributions

Conceptualisation: MAI and GFC. Data curation: MAI, NBK, AMB, FB, BN, and GFC. Formal analysis: MAI. Funding acquisition: GFC. Investigation: MAI. Methodology: MAI and GFC. Project administration: NBK and GFC. Resources: NBK and GFC. Software: GFC. Supervision: GFC. Validation: MAI. Visualisation: MAI. Writing— original draft: MAI. Writing—review & editing: MAI, NBK, AMB, FB, BN, and GFC.

## Conflicts of interest

The authors declare no conflicts of interest.

## Funding

Grants from the Wellcome Trust Institutional Strategic Support Fund (204826/Z/16/Z), NDPH Pump Priming Fund, John Fell Fund, Robertson Foundation Fellowship, and UKRI EPSRC Award (EP/X021793/1) were awarded to GFC. For the purpose of Open Access, the author has applied a CC-BY public copyright licence to any Author Accepted Manuscript version arising from this submission.

## Notes

### Competing Interest Statement

The authors have declared no competing interest.

