## Supporting Information for "Ecological niche stability of *Biomphalaria* intermediate hosts for *Schistosoma mansoni* under extreme flooding and seasonal change"

##### S1 Text: Covariate definitions and collection procedures.

###### Spatiotemporal information

For the temporal aspect, we considered the collection **timepoint** (2022, 2023a, 2023b, and 2024) as a categorical variable, with 2022 used as the reference category. Each timepoint exhibited different environmental conditions, namely in 2022 there was extreme flooding, 2023a was dry, 2023b was rainy, and 2024 was dry following extended rains season. We also included the snail sampling **start time** as a categorical variable. This was split into: early morning (08:00–10:00), late morning (10:00–12:00), and afternoon (12:00 onwards).

For the spatial aspect, we considered the **district** (Buliisa, Pakwach, and Mayuge). For the *B. sudanica* outcome, 'Mayuge' was taken as the reference category to distinguish between the eastern and western districts. Additionally, we included our constructed **polygons** as a static spatial unit to deal with spatial autocorrelation. Lastly, using Android's Fused Location Provider through the ODK Collect application, we collected the GPS positions of the water sites. This included the **altitude**, i.e. the height above sea level.

###### Site characteristics

General site characteristics we collected include the site **landform**, where the following options were available for selection: lake beach, lake marsh, pond/swamp, rice paddy, or river/river marsh. This was treated as a categorical variable. Water depth was also recorded using a measuring stick of 2 m length. Starting from the shoreline, the **shallow end**, defined as the closest distance where sampling occurred, and the **deepest end**, the farthest distance for sampling, were measured to the nearest centimetre. Information on the **water speed** (stagnant, slow, or fast) was recorded from visual inspection and was also treated as a categorical variable. We indicated if the site was used as a **landing site** (binary variable) and the **number of boats** present was included as a numerical variable. Both variables were recorded from visual inspection, and only the boats that were currently seen in the sampling area were counted.

##### Physicochemical parameters

We recorded five parameters relating to the water quality at the sites. The **water turbidity** was measured by visual inspection after collecting water in a clear container and holding it up to natural light. The categories to select from were: low (clear water, light passes easily), medium (murky/cloudy water, little light comes through), or high (opaque water, no light passes). Using a multiparameter portable meter (HI-9813-51 pH/EC/TDS and temperature Portable Meter) we recorded the **water pH**, indicating the acidity/alkalinity of the water, with a 0.1 resolution; the **water temperature** to the nearest 0.1°C; the water's ability to pass an electrical current, i.e. the **conductivity**, in mS/cm with 0.01 resolution; and the **total dissolved solids**, indicating the total amount of inorganic and organic materials dissolved in the water, to the closest ppm (mg/L).

##### Environmental and ecological attributes

Attributes relating to the environment and ecology of the water sites were further split into five categories: environmental modifications, ground substrate, vegetation, livestock, and fish. All were recorded as binary variables indicating presence/absence. Environmental modifications included **mining** and **vegetation clearance**. The options for ground substrate included **clay (slit)**, **mud**, **plastic** (including other garbage), **rock (loose)**, **rock (solid)**, **sand**, and **shells**. Types of vegetation considered were ***Ceratophyllum***, **hyacinth**, **Nile cabbage**, ***Vallisneria***, and **water lilies**. Presence of **chickens**, **cows**, **dogs**, **ducks**, **goats**, and **pigs** was observed, along with types of fish including ***Clarias***, **electric catfish**, **Nile perch**, and **tilapia**. The information was collected based on observation at the time of collection and knowledge of presence (for livestock and fish) from the village assistant.

##### Human activities

An indication regarding human activities being performed at the water sites was recorded by visual inspection and based on the village helper's knowledge. The activities included

**domestic** (bathing, retrieving drinking/cooking/bathing water, and washing of jerry cans or other items), **recreational** (swimming and children playing in water), **fishing** (visiting the site for fishing-related activities), and **sanitation** (defecation/urination or disposal of excreta occurring at the site).

#### S2 Text: Village polygons.

To create village-specific polygons, the household locations were considered and a polygon for each individual household was generated. The polygons were then merged based on the village reported by the household head. Neighbouring households with different villages reported could lead to multiple polygons per village after merging. Therefore, the approach was adapted to identify when a village was split into more than one polygon. First, we identified internal polygons (i.e. polygons that were surrounded by another polygon) and reassigned the households located within these to the surrounding village polygon. Then, we identified polygons that consisted of $\leq 5$ households—we refer to these as extra polygons—and identified their neighbouring polygons. Households within the extra polygons were then reassigned to the neighbouring polygon with which the extra polygon in question shared the most vertices. The household polygons were merged according to their new village assignments, and the village polygon each site was located in was determined to create a new village identifier for the sites.

#### S3 Tables: Summary of total snails and shedding snails found.

**Table S3.1: Number and percentage of sites with any snails present.**

| | **2022** (Flooded) | **2023a** (Dry) | **2023b** (Rainy) | **2024** (Dry following extended rains) |
|---|---|---|---|---|
| **Buliisa** | 41 (69.5%) | 54 (88.5%) | 53 (82.8%) | 59 (86.8%) |
| **Pakwach** | 34 (85.0%) | 57 (86.4%) | 56 (90.3%) | 47 (62.9%) |
| **Mayuge** | 36 (81.8%) | 45 (90.0%) | 42 (85.7%) | 39 (95.9%) |

**Table S3.2: Number and percentage of sites with any shedding snails present.**

| | **2022** (Flooded) | **2023a** (Dry) | **2023b** (Rainy) | **2024** (Dry following extended rains) |
|---|---|---|---|---|
| **Buliisa** | 7 (11.9%) | 20 (32.8%) | 20 (31.2%) | 12 (17.6%) |
| **Pakwach** | 2 (5.0%) | 19 (28.8%) | 16 (25.8%) | 6 (9.68%) |
| **Mayuge** | 3 (6.82%) | 8 (16.0%) | 7 (14.3%) | 10 (20.4%) |

#### S4 Figures: Snails and shedding snails species breakdown per timepoint and per district.

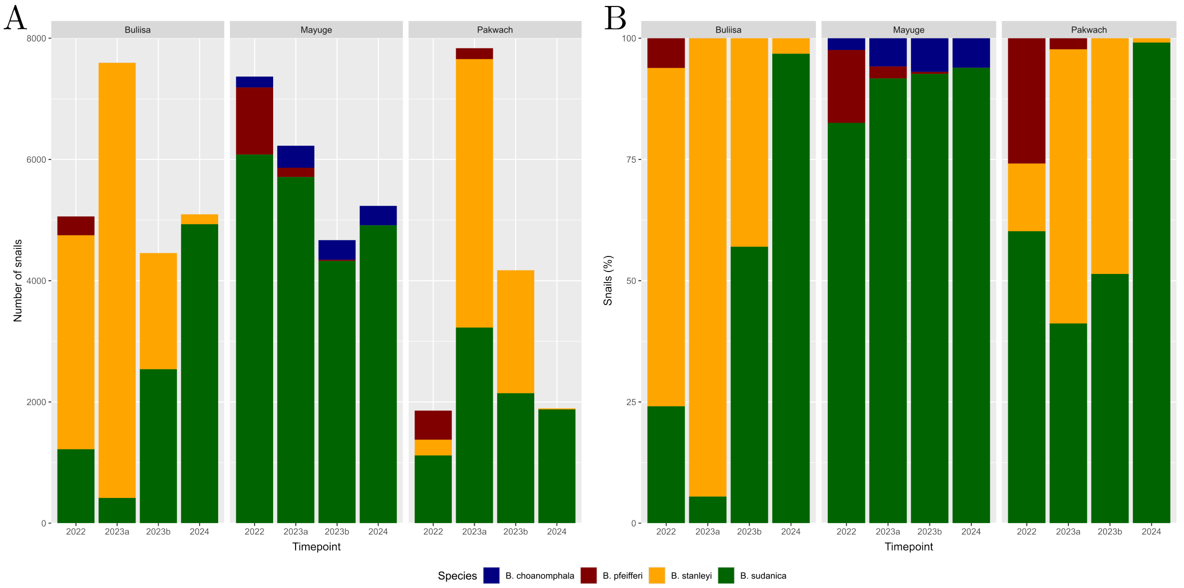


**Figure S4.1: Snails per species.** Panel A shows the total number of snails (for each species) collected at each timepoint in each district. Panel B shows the percentage of each species from the total number of snails collected at a given timepoint in a given district.

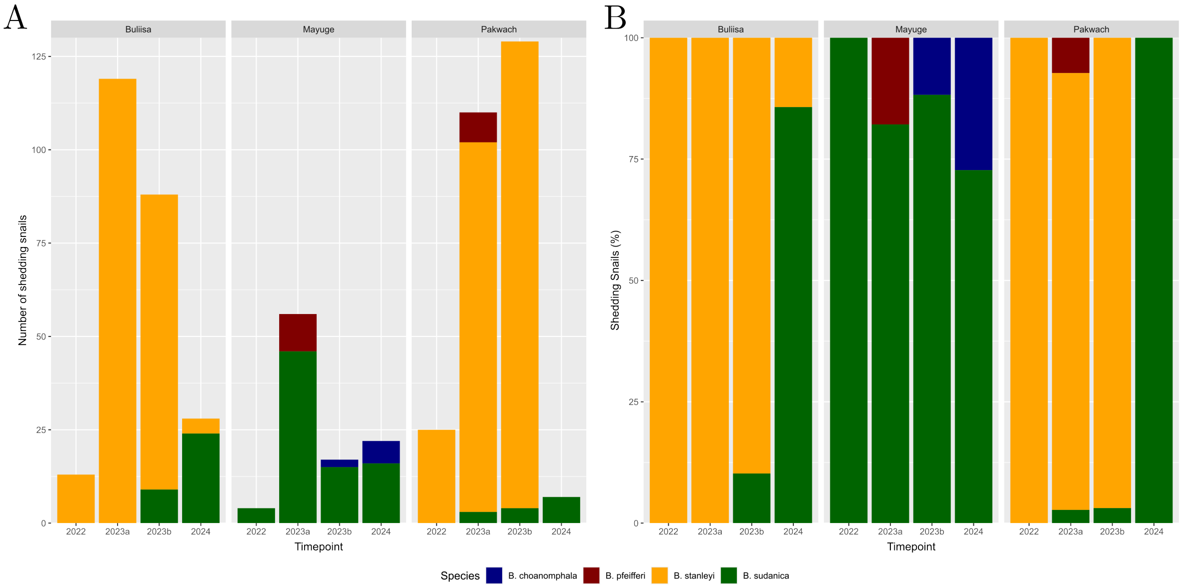


**Figure S4.2: Shedding snails per species.** Panel A shows the total number of shedding snails (for each species) collected at each timepoint in each district. Panel B shows the percentage of each species from the total number of shedding snails collected at a given timepoint in a given district.

### S5 Tables: Summary statistics of covariates exhibiting spatial and/or seasonal variation.

Table S5.1: Summary statistics at the district level.

| | Median [Q1, Q3] or Total [Proportion] | | | Difference test $p$-value | | |
|---|---|---|---|---|---|---|
| Variable | Buliisa ($N = 252$) | Pakwach ($N = 230$) | Mayuge ($N = 192$) | Buliisa/ Pakwach | Buliisa/ Mayuge | Pakwach/ Mayuge |
| **Spatiotemporal** | | | | | | |
| altitude | 621 [612, 631] | 628 [618, 637] | 1149 [1139, 1159] | 0.004 | <0.001 | <0.001 |
| **Site characteristics** | | | | | | |
| landform | | | | | | |
| lake beach | 126 [0.500] | 27 [0.117] | 67 [0.349] | <0.001 | 0.006 | <0.001 |
| lake marsh | 92 [0.365] | 95 [0.413] | 71 [0.370] | 0.970 | 1.000 | 1.000 |
| pond/swamp | 15 [0.060] | 33 [0.143] | 40 [0.208] | 0.010 | <0.001 | 0.310 |
| rice paddy | 1 [0.004] | 2 [0.009] | 12 [0.063] | 1.000 | 0.003 | 0.015 |
| river/river marsh | 18 [0.071] | 73 [0.317] | 2 [0.010] | <0.001 | 0.014 | <0.001 |
| shallow end | 22 [14, 29] | 21 [14, 30] | 16.5 [12, 23] | 1.000 | <0.001 | <0.001 |
| deepest end | 69 [54.8, 86] | 79 [54, 91] | 60 [39.5, 84.2] | 0.028 | 0.004 | <0.001 |
| water speed | | | | | | |
| stagnant | 59 [0.234] | 105 [0.457] | 79 [0.411] | <0.001 | <0.001 | 1.000 |
| slow | 111 [0.440] | 98 [0.426] | 64 [0.333] | 1.000 | 0.085 | 0.193 |
| fast | 82 [0.325] | 27 [0.117] | 49 [0.255] | <0.001 | 0.400 | 0.001 |
| landing site | 99 [0.39] | 65 [0.28] | 37 [0.19] | 0.042 | <0.001 | 0.126 |
| **Physicochemical** | | | | | | |
| water turbidity | | | | | | |
| low | 92 [0.365] | 84 [0.365] | 73 [0.380] | 1.000 | 1.000 | 1.000 |
| medium | 137 [0.544] | 104 [0.452] | 91 [0.474] | 0.170 | 0.520 | 1.000 |
| high | 23 [0.091] | 42 [0.183] | 28 [0.146] | 0.015 | 0.305 | 1.000 |
| water pH | 8.6 [7.7, 9] | 7.6 [7.4, 8.2] | 7.6 [7.1, 8.4] | <0.001 | <0.001 | 0.385 |
| water temperature | 27.7 [26.3, 29] | 29.2 [27.8, 30.8] | 28.6 [27, 30.2] | <0.001 | <0.001 | 0.001 |
| conductivity | 0.48 [0.41, 0.56] | 0.37 [0.25, 0.47] | 0.11 [0.08, 0.27] | <0.001 | <0.001 | 0.001 |
| **Environmental and ecological** | | | | | | |
| vegetation clearance | 23 [0.091] | 45 [0.196] | 77 [0.401] | 0.005 | <0.001 | <0.001 |
| mud | 122 [0.484] | 159 [0.691] | 108 [0.562] | <0.001 | 0.370 | 0.025 |
| plastic | 125 [0.496] | 77 [0.335] | 69 [0.359] | 0.001 | 0.016 | 1.000 |
| rock loose | 23 [0.091] | 3 [0.013] | 32 [0.167] | <0.001 | 0.075 | <0.001 |
| rock solid | 29 [0.115] | 7 [0.030] | 45 [0.234] | 0.002 | 0.004 | <0.001 |
| sand | 193 [0.766] | 125 [0.543] | 104 [0.542] | <0.001 | <0.001 | 1.000 |
| *Ceratophyllum* | 71 [0.282] | 72 [0.313] | 16 [0.083] | 1.000 | <0.001 | <0.001 |
| hyacinth | 173[0.687] | 205 [0.897] | 149 [0.776] | <0.001 | 0.141 | 0.006 |
| *Vallisneria* | 112 [0.444] | 48 [0.209] | 33 [0.172] | <0.001 | <0.001 | 1.000 |
| chickens | 67 [0.266] | 9 [0.039] | 25 [0.130] | <0.001 | 0.002 | 0.004 |
| dogs | 78 [0.310] | 48 [0.209] | 41 [0.214] | 0.048 | 0.094 | 1.000 |
| goats | 150 [0.595] | 117 [0.509] | 124 [0.646] | 0.207 | 0.969 | 0.019 |
| pigs | 88 [0.349] | 56 [0.243] | 55 [0.286] | 0.045 | 0.582 | 1.000 |
| electric catfish | 18 [0.071] | 26 [0.113] | 0 [0.000] | 0.4615 | <0.001 | <0.001 |
| Nile perch | 190 [0.75] | 117 [0.61] | 153 [0.67] | 0.122 | 0.005 | 0.830 |
| **Human activities** | | | | | | |
| sanitation | 130 [0.516] | 63 [0.274] | 62 [0.323] | <0.001 | <0.001 | 0.965 |

#### Table S5.2: Summary statistics per timepoint.

| | Median [Q1, Q3] or Total [Proportion] | | | | Difference test $p$-value | | | | | |
|---|---|---|---|---|---|---|---|---|---|---|
| **Variable** | **2022** ($N = 143$) | **2023a** ($N = 177$) | **2023b** ($N = 175$) | **2024** ($N = 179$) | **2022/ 2023a** | **2022/ 2023b** | **2022/ 2024** | **2023a/ 2023b** | **2023a/ 2024** | **2023b/ 2024** |
| **Site characteristics** | | | | | | | | | | |
| landform | | | | | | | | | | |
| lake beach | 55 | 80 | 54 | 31 | 1.000 | 1.000 | <0.001 | 0.047 | <0.001 | 0.026 |
| lake marsh | 46 | 44 | 69 | 99 | 1.000 | 1.000 | <0.001 | 0.029 | <0.001 | 0.024 |
| pond/swamp | 28 | 25 | 23 | 12 | 1.000 | 0.964 | 0.006 | 1.000 | 0.204 | 0.385 |
| rice paddy | 2 | 4 | 5 | 4 | 1.000 | 1.000 | 1.000 | 1.000 | 1.000 | 1.000 |
| river/river marsh | 12 | 24 | 24 | 33 | 1.000 | 1.000 | <0.093 | 1.000 | 1.000 | 1.000 |
| shallow end | 18 [13, 27] | 20 [12, 27] | 18 [10, 27] | 22 [18, 27.5] | 1.000 | 0.227 | 0.169 | 0.533 | 0.032 | <0.001 |
| deepest end | 69 [53.5, 90.5] | 70 [52, 85] | 53 [40, 79.5] | 79 [66, 92.5] | 0.553 | <0.001 | 0.007 | 0.002 | <0.001 | <0.001 |
| landing site | 37 [0.26] | 39 [0.22] | 79 [0.45] | 46 [0.26] | 1.000 | 0.004 | 1.000 | <0.001 | 1.000 | 0.001 |
| **Physicochemical** | | | | | | | | | | |
| water pH | 8.4 [7.6, 9] | 8.3 [7.5, 9] | 8 [7.4, 8.7] | 7.5 [7.2, 8] | 1.000 | 0.054 | <0.001 | 0.115 | <0.001 | <0.001 |
| water temperature | 28.1 [26, 29.5] | 28.8 [27.4, 30] | 29.7 [28.2, 31.5] | 27.4 [26.3, 28.6] | 0.003 | <0.001 | 0.139 | <0.001 | <0.001 | <0.001 |
| conductivity | 0.43 [0.32, 0.58] | 0.43 [0.22, 0.48] | 0.27 [0.14, 0.40] | 0.46 [0.22, 0.54] | 0.145 | <0.001 | 1.000 | <0.001 | 0.563 | <0.001 |
| **Environmental and ecological** | | | | | | | | | | |
| vegetation clearance | 21 [0.147] | 35 [0.198] | 49 [0.280] | 40 [0.223] | 1.000 | 0.040 | 0.660 | 0.550 | 1.000 | 1.000 |
| clay | 5 [0.035] | 16 [0.090] | 17 [0.097] | 44 [0.246] | 0.467 | 0.306 | <0.001 | 1.000 | <0.001 | 0.002 |
| mud | 75 [0.524] | 88 [0.497] | 104 [0.594] | 122 [0.682] | 1.000 | 1.000 | 0.035 | 0.510 | 0.004 | 0.660 |
| plastic | 53 [0.371] | 39 [0.220] | 89 [0.509] | 90 [0.503] | 0.028 | 0.113 | 0.143 | <0.001 | <0.001 | 1.000 |
| rock loose | 26 [0.182] | 8 [0.045] | 10 [0.057] | 14 [0.078] | 0.001 | 0.006 | 0.051 | 1.000 | 1.000 | 1.000 |
| shells | 11 [0.077] | 29 [0.164] | 28 [0.160] | 85 [0.475] | 0.180 | 0.230 | <0.001 | 1.000 | <0.001 | <0.001 |
| *Ceratophyllum* | 21 [0.147] | 36 [0.203] | 38 [0.217] | 64 [0.358] | 1.000 | 0.867 | <0.001 | 1.000 | 0.011 | 0.031 |
| hyacinth | 86 [0.601] | 122 [0.689] | 151 [0.863] | 168 [0.939] | 0.770 | <0.001 | <0.001 | <0.001 | <0.001 | 0.164 |
| Nile cabbage | 43 [0.30] | 30 [0.17] | 48 [0.27] | 13 [0.07] | 0.049 | 1.000 | <0.001 | 0.151 | 0.050 | <0.001 |
| *Vallisneria* | 36 [0.252] | 75 [0.424] | 66 [0.377] | 16 [0.089] | 0.012 | 0.142 | <0.001 | 1.000 | <0.001 | <0.001 |
| water lilies | 15 [0.105] | 11 [0.062] | 45 [0.257] | 2 [0.011] | 1.000 | 0.006 | 0.003 | 0.003 | 0.135 | <0.001 |
| chickens | 28 [0.196] | 6 [0.034] | 41 [0.234] | 26 [0.145] | <0.001 | 1.000 | 1.000 | <0.001 | 0.003 | 0.272 |
| cows | 106 [0.741] | 100 [0.565] | 133 [0.760] | 116 [0.648] | 0.010 | 1.000 | 0.564 | 0.001 | 0.808 | 0.171 |
| dogs | 16 [0.112] | 6 [0.034] | 74 [0.423] | 71 [0.397] | 0.071 | <0.001 | <0.001 | <0.001 | <0.001 | 1.000 |
| goats | 75 [0.524] | 78 [0.441] | 127 [0.726] | 111 [0.620] | 1.000 | 0.002 | 0.641 | <0.001 | 0.006 | 0.271 |
| pigs | 40 [0.280] | 16 [0.090] | 57 [0.326] | 86 [0.480] | <0.001 | 1.000 | 0.002 | <0.001 | <0.001 | 0.026 |
| *Clarias* | 37 [0.259] | 70 [0.395] | 86 [0.491] | 120 [0.670] | 0.084 | <0.001 | <0.001 | 0.530 | <0.001 | 0.006 |
| electric catfish | 4 [0.028] | 6 [0.034] | 20 [0.114] | 14 [0.078] | 1.000 | 0.043 | 0.528 | 0.044 | 0.677 | 1.000 |
| Nile perch | 78 [0.55] | 106 [0.60] | 141 [0.81] | 135 [0.75] | 1.000 | <0.001 | <0.001 | <0.001 | 0.015 | 1.000 |

### S6 Figures: Distributions of key covariates.

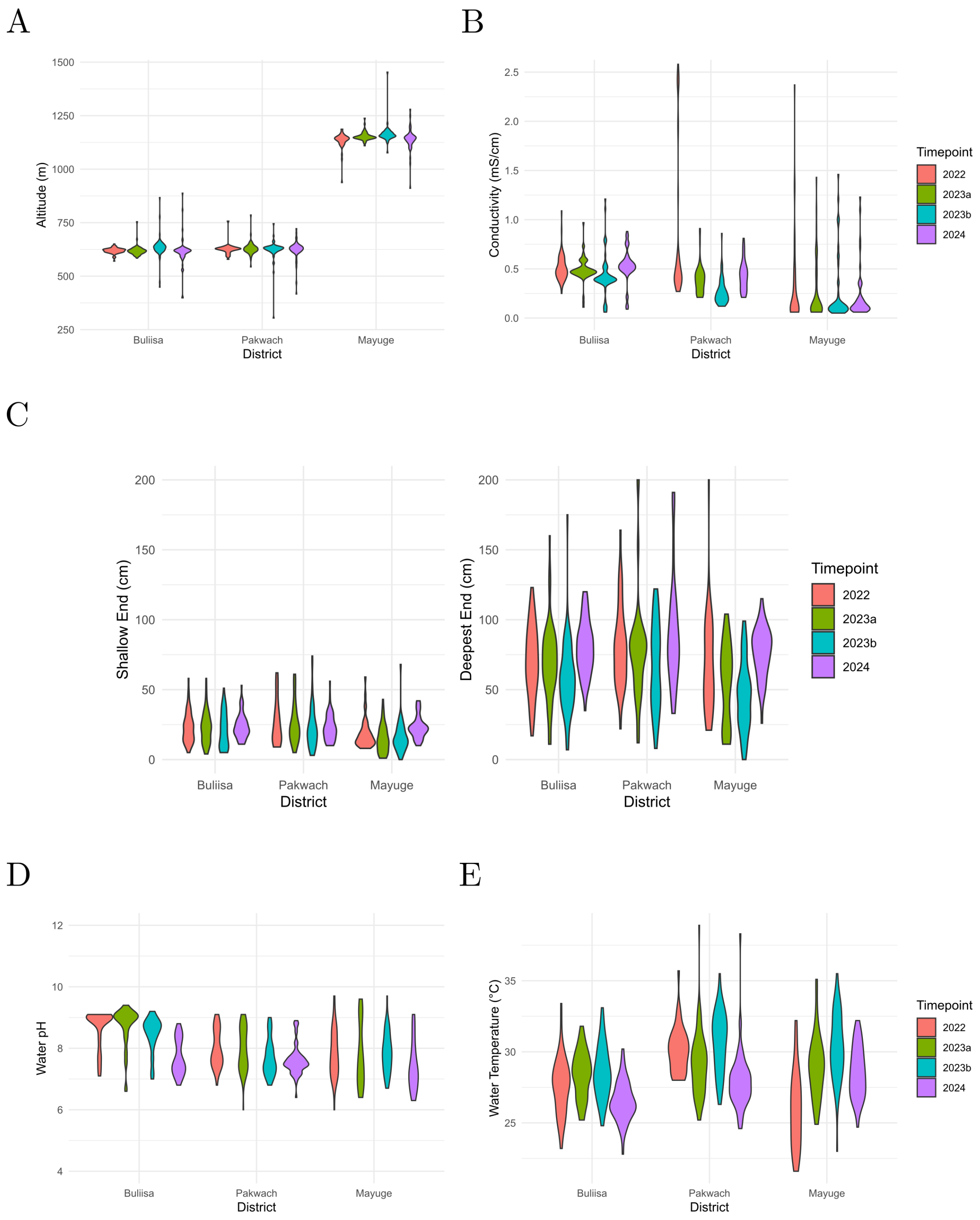


**Figure S6.1: Distributions of key covariates.** A. altitude, B. conductivity, C. Water depth (left: shallow end; right: deepest end), D. water pH, E. water temperature.

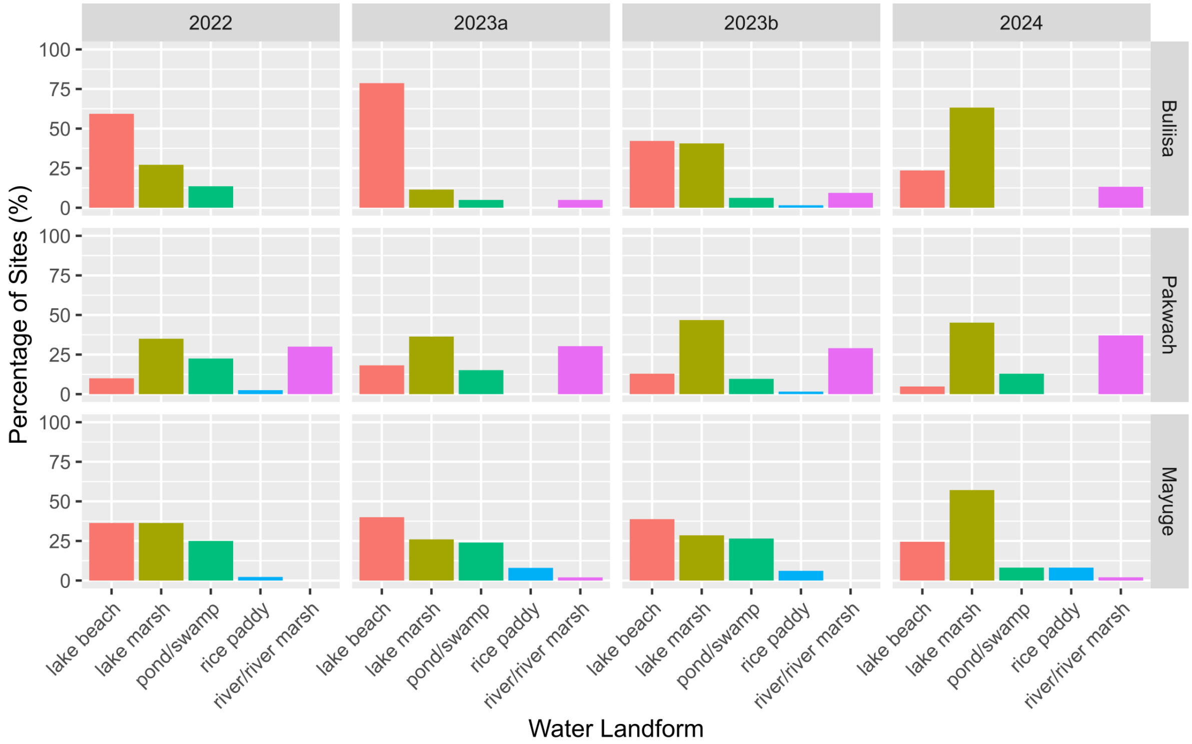


**Figure S6.2: Percentage of sites with each landform (per timepoint, per district).**

#### S7 Figure: Optimal number of clusters based on the gap statistic.

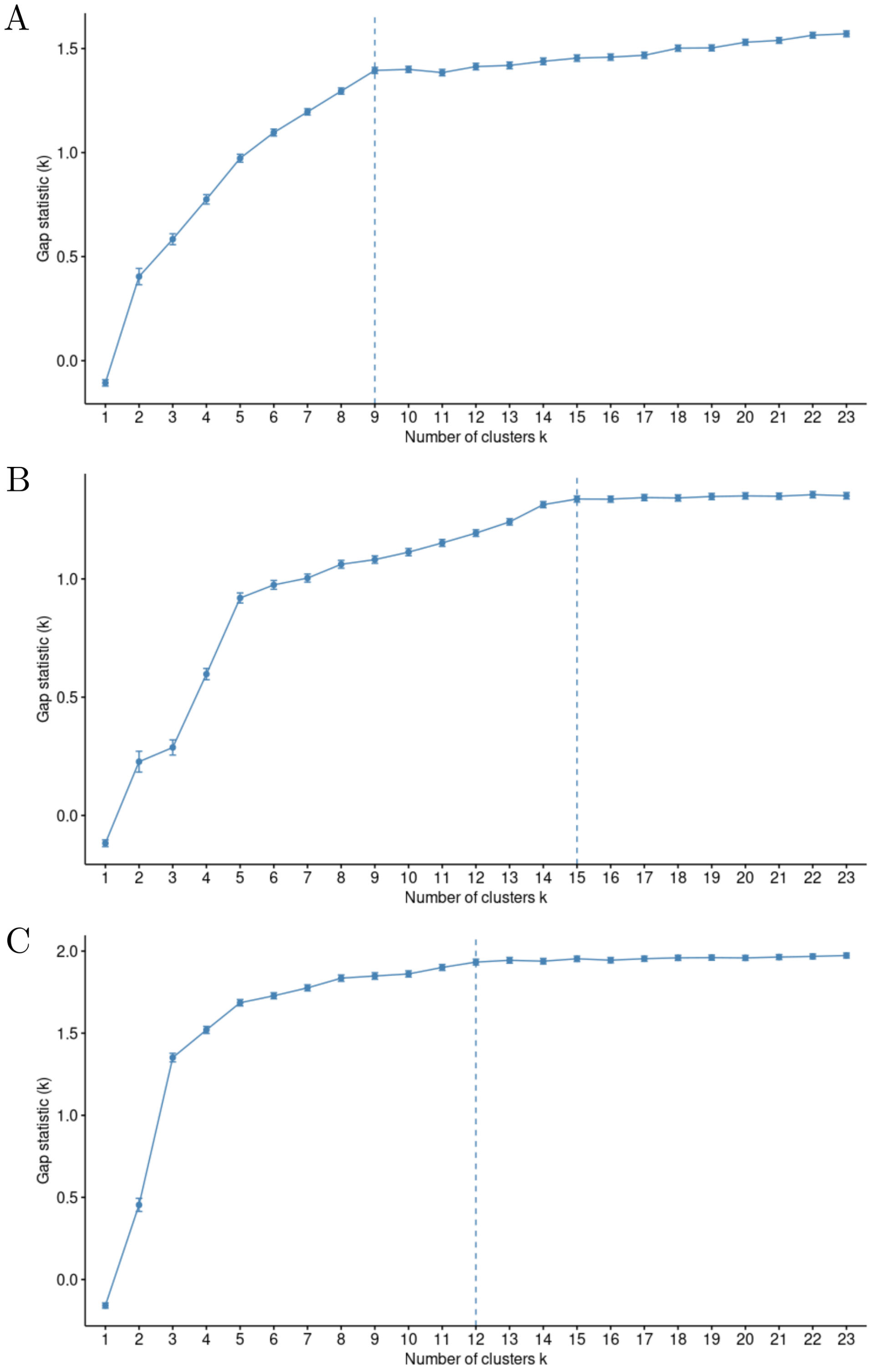


**Figure S7.1: Optimal number of clusters based on the gap statistic.** A. Buliisa, B. Pakwach, C. Mayuge.

#### S8 Figures: Village-specific polygons.

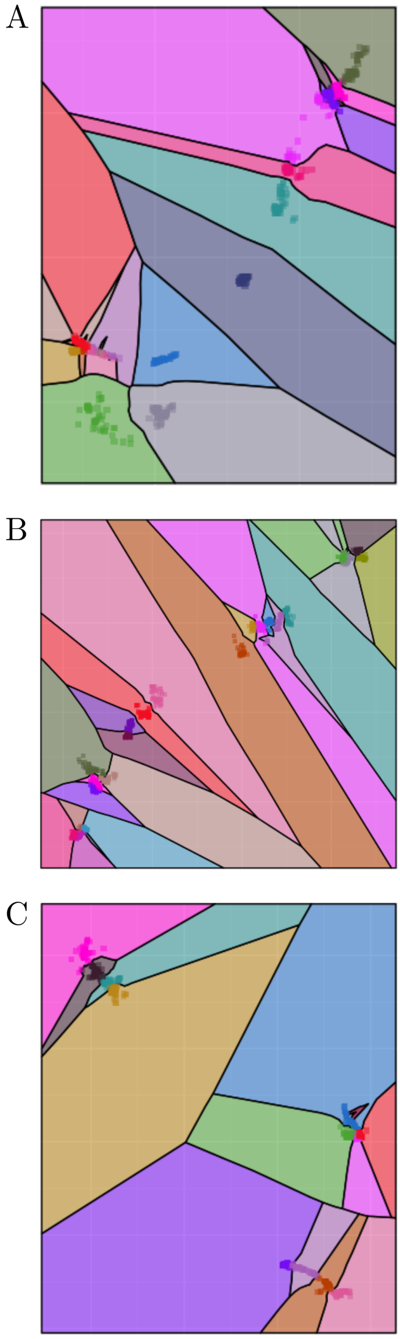


**Figure S8.1: Village-specific polygons** Merged Voronoi polygons using household locations and self-reported villages. A. Buliisa, B. Pakwach, C. Mayuge. Squares represent each household.

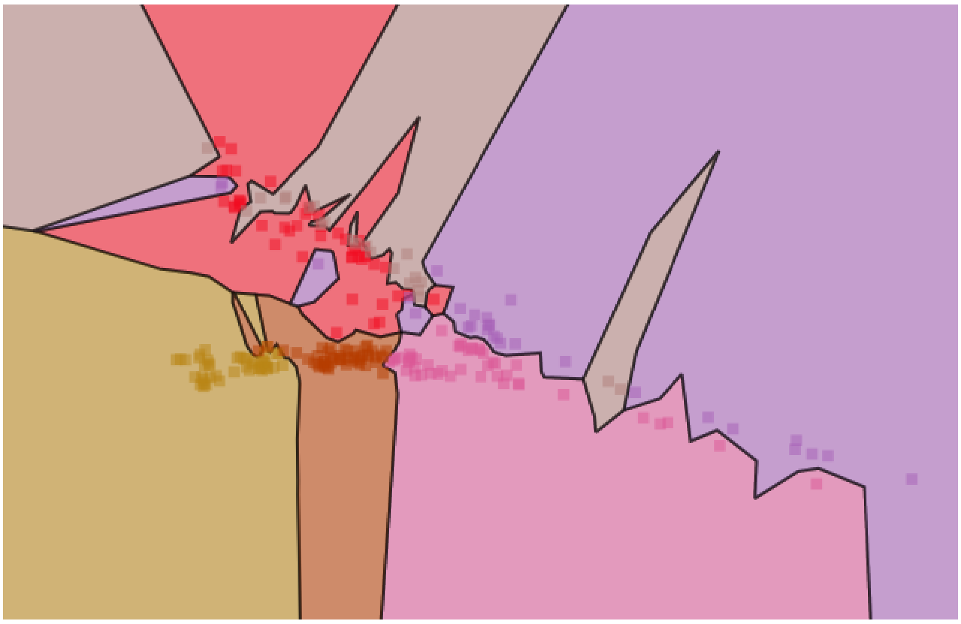

**Figure S8.2:** Close-up of panel A from Figure S8.1 where villages overlap and are represented by multiple distinct polygons.

#### S9 Figure: Fully adjusted *B. sudanica* model excluding Mayuge.

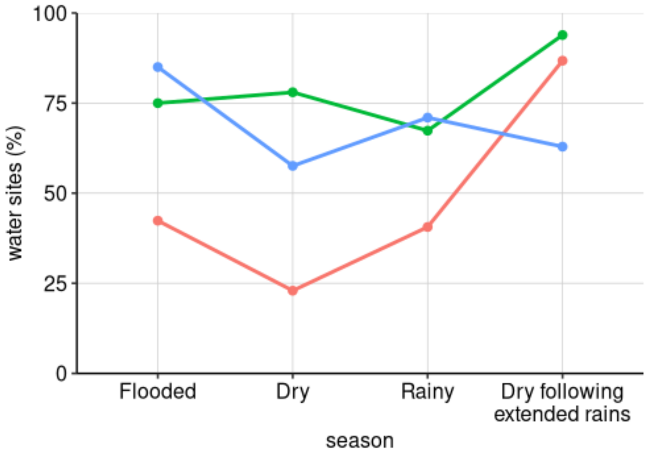


**Figure S9.1: Water sites with *B. sudanica* present per district and timepoint.** The numbers of sites with presence of this species are as follows in order of season presented: Buliisa (red): 25/59, 14/61, 26/64, 59/68; Pakwach (blue): 34/40, 38/66, 44/62, 39/62; Mayuge (green): 33/44, 39/50, 33/49, 46/49.

#### S10 Figure: Fully adjusted *B. sudanica* model excluding Mayuge.

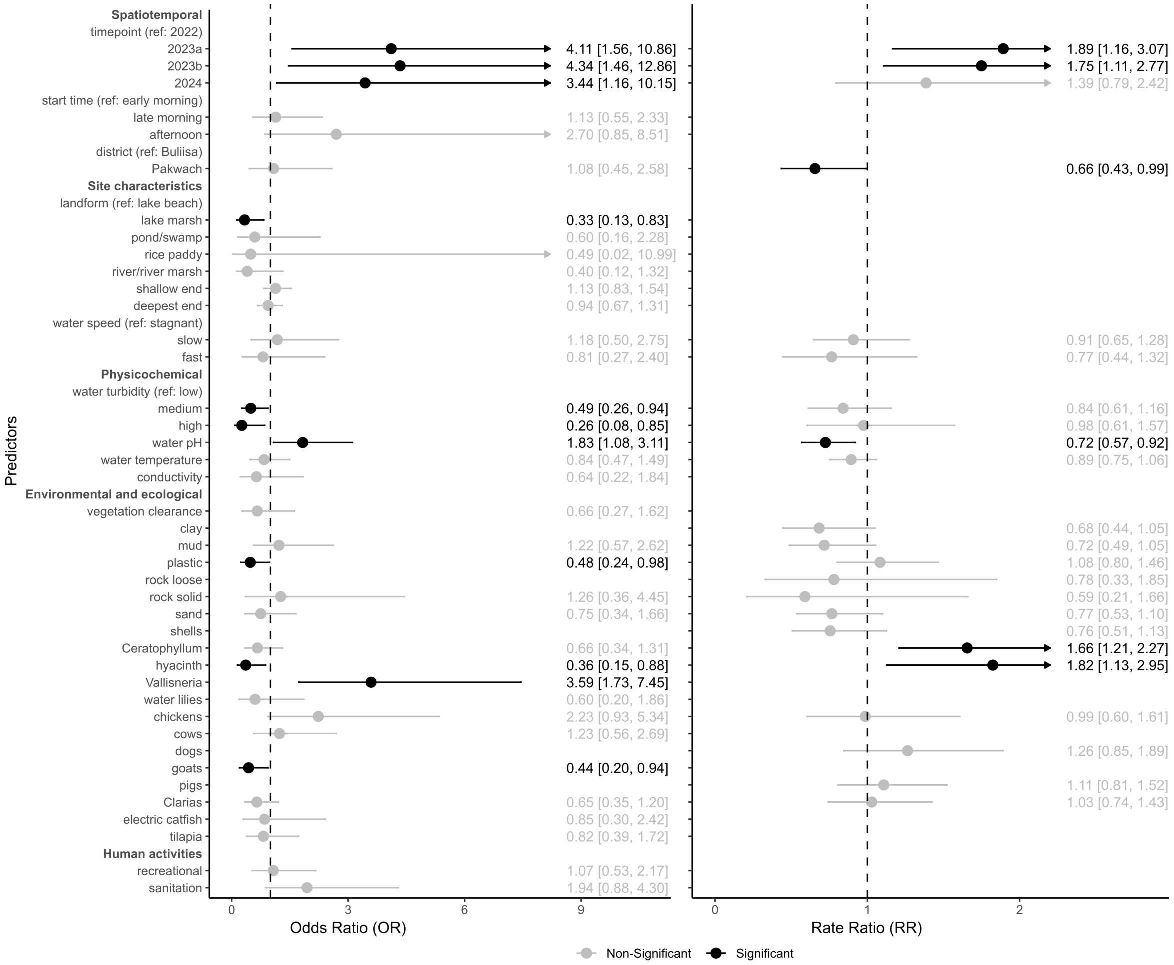


**Figure S10.1:** Odds Ratios (left) and Rate Ratios (right) for the zero-inflation and conditional components of the model, respectively. Dots correspond to the exponentiated coefficient estimates and lines correspond to the 95% confidence intervals. Arrows indicate that the confidence interval continues off the plot.

#### S11 Figure: Fully adjusted *B. sudanica* model excluding Mayuge.

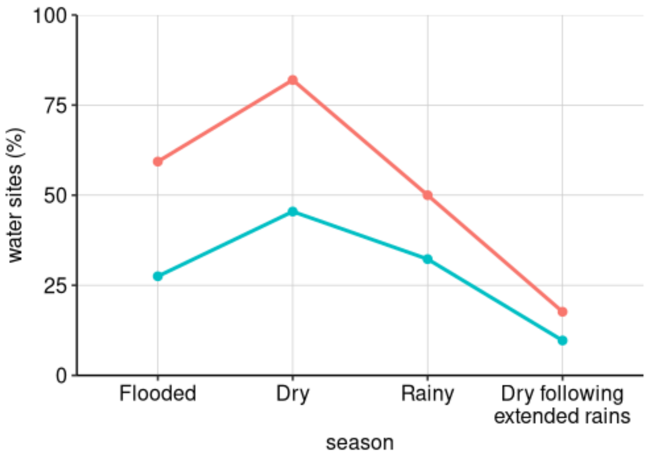


**Figure S11.1: Water sites with *B. stanleyi* present per district and timepoint.** The numbers of sites with presence of this species are as follows in order of season presented: Buliisa (red): 35/59, 50/61, 32/64, 12/68; Pakwach (teal): 311/40, 30/66, 20/62, 6/62.

#### S10 Figure: Residual goodness-of-fit tests.

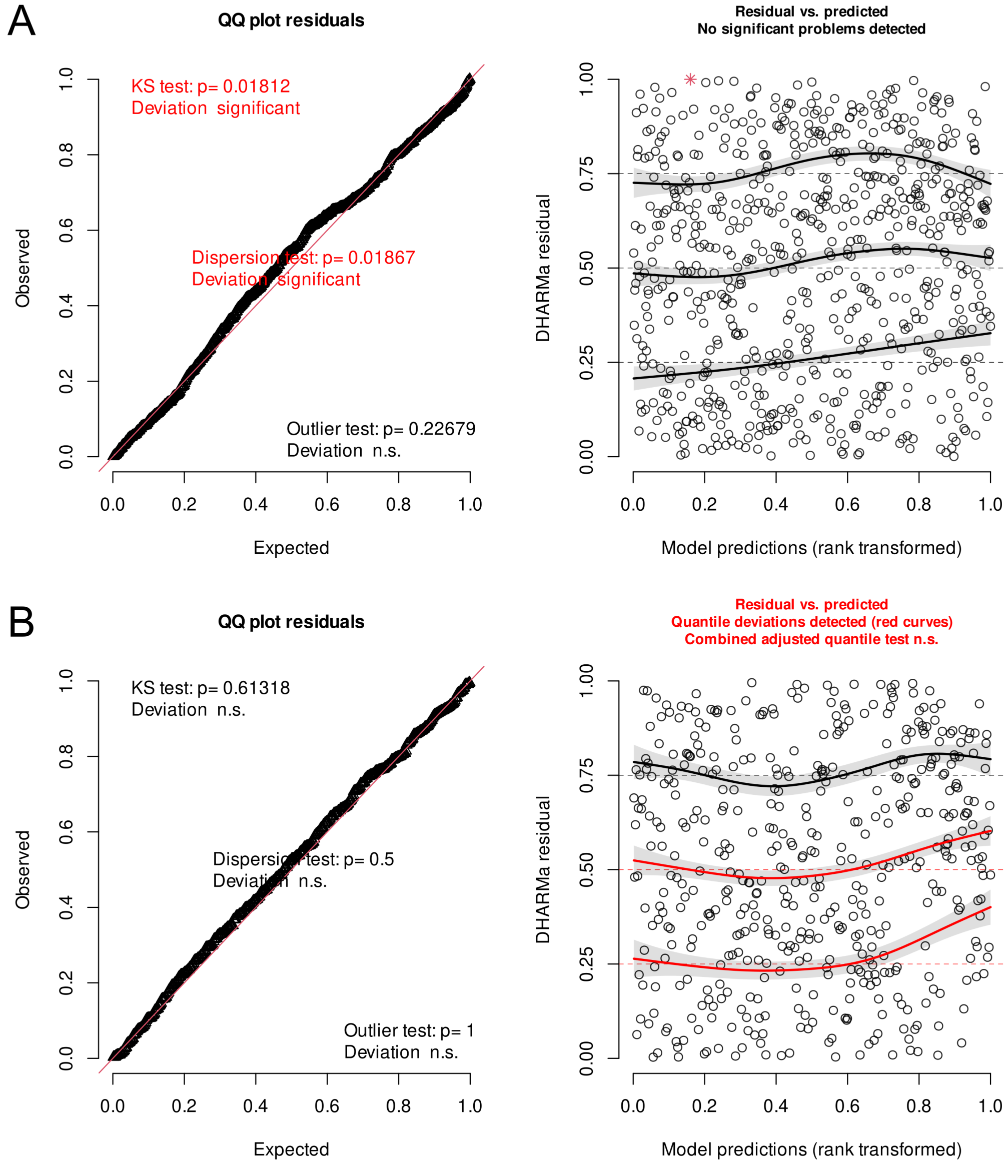


**Figure S10.1: Residual goodness-of-fit tests.** A. Residuals for *B. sudanica* model, B. Residuals for *B. stanleyi* model.
